# Chromatin loading of MCM hexamers is associated with di-/tri-methylation of histone H4K20 toward S phase entry

**DOI:** 10.1101/2021.05.14.443718

**Authors:** Yoko Hayashi-Takanaka, Yuichiro Hayashi, Yasuhiro Hirano, Atsuko Miyawaki-Kuwakado, Yasuyuki Ohkawa, Chikashi Obuse, Hiroshi Kimura, Tokuko Haraguchi, Yasushi Hiraoka

## Abstract

DNA replication is a key step in initiating cell proliferation. Loading hexameric complexes of minichromosome maintenance (MCM) helicase onto DNA replication origins during the G1 phase is essential for initiating DNA replication. Here, we examined MCM hexamer states during the cell cycle in human hTERT-RPE1 cells using multicolor immunofluorescence-based, single-cell plot analysis, and biochemical size fractionation. Experiments involving cell-cycle arrest at the G1 phase and release from the arrest revealed that a double MCM hexamer was formed via a single hexamer during G1 progression. A single MCM hexamer was recruited to chromatin in the early G1 phase. Another single hexamer was recruited to form a double hexamer in the late G1 phase. We further examined relationship between the MCM hexamer states and the methylation levels at lysine 20 of histone H4 (H4K20) and found that the double MCM hexamer state was correlated with di/trimethyl-H4K20 (H4K20me2/3). Inhibiting the conversion from monomethyl-H4K20 (H4K20me1) to H4K20me2/3 retained the cells in the single MCM hexamer state. Non-proliferative cells, including confluent cells or Cdk4/6 inhibitor-treated cells, also remained halted in the single MCM hexamer state. We propose that the single MCM hexamer state is a halting step in the determination of cell cycle progression.

## Introduction

Replication of genomic DNA is crucial for the proliferation of living organisms. The G1 phase of the cell cycle is an important step in the preparation of DNA replication in the S phase (1,2). During the G1 phase, mammalian cells decide either to enter the S phase for proliferation or halt at the G1 phase for quiescence (3). While most cancer cells lose the ability to be arrested at the G1 phase, normal cells are typically arrested in the G1 phase in response to the accumulation of DNA damage or external stimuli, resulting in a prolonged cell cycle (4). Once the cells pass through the G1 checkpoint, they enter the S phase within 2–3 h, with no need for other stimuli (5). Therefore, the decision to exit the G1 phase is a key event in entering the S phase for cell proliferation.

Prior to the S phase, the assembly of the “pre-replication complex” (pre-RC) at the DNA replication origins on chromatin is an essential step during the G1 phase ((6,7) reviewed in (8-10)). This complex consists of the origin recognition complex (ORC), Cdc6, Cdt1, and minichromosome maintenance protein (MCM). By the end of the G1 phase, two copies of the MCM hexamer (Mcm2-7) are loaded on the replication origin to function as a DNA helicase in the S phase. Metazoan replication origins do not share a clear consensus sequence, although they tend to have enriched GC-rich sequences and G4 quadruplex structures (11). In contrast, in the budding yeast *Saccharomyces cerevisiae*, an autonomously replicating sequence containing the replication origin has been previously identified (12,13). *In vitro* studies using recombinant yeast proteins found that Orc1-6, Cdc6, Cdt1, and Mcm2-7 are sufficient for the pre-RC assembly of the replication origin (14,15). Two hexamer complexes of MCM are sequentially loaded on chromatin: first, a single MCM hexamer is recruited to the replication origin where the ORC complex has already been loaded with the help of Cdc6 and Cdt1 (16,17); then, a second MCM hexamer is loaded to form a double MCM hexamer. Structures of the single and double MCM hexamers and their intermediates were also determined using the *in vitro* reconstituted complexes of recombinant yeast proteins by cryo-electron microscopy (18,19). In mammalian cells, however, it is still unknown when and how single and double MCM hexamers are loaded *in vivo* on chromatin in the G1 phase. In addition, the chromatin state required for MCM hexamer formation remains to be elucidated.

Histone modification is also involved in replication initiation (20). Lysine 20 of histone H4 (H4K20) is a major methylated residue on the histone H4 tail that can exist in mono-, di-, and tri-methylation forms (H4K20me1, H4K20me2, and H4K20me3, respectively). H4K20 methylation is involved in DNA replication, including the initiation step and maintenance of genome integrity (21). H4K20me2 and H4K20me3, which are methylated from H4K20me1 by Suv420h1 and Suv420h2 (also called KMT5B and KMT5C, respectively), serve as binding sites for mammalian Orc1 and its cofactors (22,23). Moreover, the histone methyltransferase Set8 (also known as KMT5A or PR-set7), which catalyzes the monomethylation of histone H4 at K20, is involved in chromatin decompaction at the M/G1 transition (24) and is required for S phase progression (25,26). It has also been reported that Set8 recruits Orc1 and MCM to chromatin *in vitro* (27). Therefore, the methylation of histone H4K20 by Set8 and/or Suv420h1/2 methyltransferase is crucial for pre-RC assembly on chromatin during the G1 phase to initiate DNA replication.

Here, we studied the MCM assembly on chromatin in relation to the histone H4K20 methylation levels during the G1 phase using multicolor immunofluorescence-based, single-cell plot analysis (hereafter, referred to as single-cell plot analysis) (28), which enables quantitative analysis in individual cells based on their fluorescence microscopic images.

## Materials and Methods

### Antibodies

The primary antibodies used for immunostaining are as follows: mouse monoclonal anti-Mcm3 (sc-390480; Santa Cruz Biotechnology), rabbit monoclonal anti-Mcm2 (D7G11; Cell Signaling Technology), rabbit monoclonal anti-Mcm2 S108 phosphorylation (EPR4121; Abcam), rabbit monoclonal anti-Cdt1 (D10F11; Cell Signaling Technology), rabbit monoclonal anti-Cdc45 (D7G6; Cell Signaling Technology), and rabbit monoclonal anti-phosphorylated Rb (Ser807/811) (D20B12; Cell Signaling Technology). The secondary antibodies used were as follows: donkey anti-mouse IgG (711-005-151; Jackson ImmunoResearch) and donkey anti-rabbit IgG (711-005-152; Jackson ImmunoResearch). Mouse monoclonal antibodies against histone H4K20me1 (CMA421), H4K20me2 (CMA422), and H4K20me3 (CMA423) were generated as previously described (29).

### Cell culture

hTERT-RPE1 (ATCC, CRL-4000), HeLa (RCB0007), U2OS (ATCC, HTB-96), MRC5 (ATCC, CCL-171), and IMR90 (ATCC, CCL-186) cell lines were grown in a culture medium comprising high-glucose Dulbecco’s modified Eagle’s medium (Sigma-Aldrich) supplemented with penicillin/streptomycin (100 units/mL penicillin, 100 μg/mL streptomycin; Fujifilm), and 10% fetal calf serum (Thermo Fisher Scientific), as described previously (30,31). Contact inhibition was achieved by incubating the confluent cells for four additional days after they covered almost the entire area of the adhesive surface; the medium was changed every day. siRNA transfection was performed using Lipofectamine RNAiMAX (Thermo Fisher Scientific) with siRNAs (SASI_Hs01_00180215 for Set8, SASI_Hs02_00348728 for Suv420h1, and Universal Negative Control #1, Sigma-Aldrich) according to the manufacturer’s protocol. Unless otherwise specified, the cells were harvested 2 days after transfection (>40 h).

### Cell synchronization

To synchronize hTERT-RPE1 cells at the G1 phase, palbociclib (Sigma-Aldrich) was used as previously described (32,33) with modifications. Cells were plated at 1.3 × 10^4^ cells/cm^2^, incubated for 6 h, and treated with 1 µM or 150 nM palbociclib for 18 h. To release the cells from the arrest, the cells that had been treated with 150 nM palbociclib were washed three times with pre-warmed culture medium, and then cultured until collection at the indicated times. To synchronize hTERT-RPE1 cells in the S phase, the cells were plated and incubated as described above, and then treated with 200 µM hydroxyurea (HU) (Fujifilm) (34) for 18 h.

### Fluorescence staining for single-cell plot analysis

Immunostaining was performed as previously described (31,35,36) with minor modifications. Cells were plated in a 12-well plate containing a coverslip (15 mm diameter; No. 1S; Matsunami) for more than 1 day. In the pre-extraction method, to extract soluble proteins, cells in the living state were first treated with 1 ml of 0.2% Triton X-100 solution containing 20 mM HEPES [pH 7.4], 100 mM NaCl, and 300 mM sucrose for 5 min on ice. After removing the Triton X-100 solution, the cells were fixed with 1 ml of the fixative (2% paraformaldehyde (Electron Microscopy Sciences) dissolved in 250 mM HEPES [pH 7.4]) for 10 min at room temperature (∼25°C). In the direct fixation method, the cells were first fixed with the fixative described above. After fixation, cells were permeabilized with 1% Triton X-100 in PBS for 15 min and blocked with Blocking One (Nacalai Tesque), as described previously (35,36). The fixed cells were incubated with the indicated primary antibodies (0.2–1 μg/mL) for 2 h at room temperature (∼25°C) and washed three times (10 min each) with PBS. The cells were treated with secondary antibodies (1 μg/mL) and 0.1 μg/mL Hoechst 33342 for 2 h at room temperature (∼25°C) and washed with PBS. The cells were mounted in Prolong Gold or Prolong Glass Antifade (Thermo Fisher Scientific).

To label replication foci, the cells were cultured in culture medium containing 10 μM 5-ethynyl-2-deoxyuridine (EdU) for 30 min, fixed with the fixative for 10 min, and the resulting EdU signals were detected with Alexa Fluor 647 using a Click-iT EdU Imaging Kit (Thermo Fisher Scientific) (29). For the pulse-chase experiments, the cells were cultured in culture medium containing 10 μM EdU for 30 min, washed three times with the pre-warmed culture medium, and then cultured with the medium not containing EdU until fixation at the indicated times.

### Microscopy

Fluorescence images were collected using a DeltaVision Elite system (GE Healthcare Inc.) equipped with a pco.edge 4.2 sCMOS camera (PCO) and an Olympus 40× UApo/340 oil immersion objective lens (NA 0.65 with iris) with DeltaVision Elite Filter Sets (DAPI-FITC-TRITC-Cy5).

### Single-cell plot analysis

Single-cell plot analysis is a method used to measure the levels of multiple intracellular components based on fluorescence microscopic images and plot their correlation in individual cells (28). The fluorescence intensity of the nuclei was measured using the NIS Elements software (version 3.0; Nikon). The background intensity outside the cells was subtracted from that of the total image, and the nuclear areas of individual cells were determined by automatic thresholding using Hoechst signals. The sum intensity (i.e., the average intensity × nuclear area) in each nucleus was measured for all the fluorescence channels. Cells in the M phase were not used for the analysis because the area of condensed chromosomes differed substantially from that of nuclei in the interphase (28). To plot the Hoechst intensity distribution of 400–450 nuclei, the signal intensity in each nucleus was normalized to the nuclei with the 10th lowest and 10th highest intensity, set as 1 and 2, respectively, because the nuclei with the lowest and highest intensities were sometimes outliers. The Hoechst intensity was plotted on a linear scale. The average intensity of EdU, Mcm2/3, phosphorylated Mcm2 at the 108^th^ serine residue (Mcm2-S108ph), phosphorylated Rb (Rb-ph), histone H4K20me1/me2/me3, Cdt1, and Cdc45 was set to 1 (0 on the log_2_ scale) and plotted on a log_2_ scale. In the siRNA experiments (Fig. 4B, 6D and Fig. S4), the fluorescence intensities were normalized to those of the control siRNA.

Values relative to the average were plotted using Mathematica (version 11 or 12) (Wolfram Research).

### Estimation of duration of each cell cycle phase

The total cell cycle length was determined from the growth curve of each cell type (Fig. 3B). The duration of each cell cycle step was calculated by multiplying the ratio of the number of cells in each cell cycle step by the total number of cells by the total cell cycle length. The number of cells in the S phase was counted by the EdU signal, and the numbers of cells in the G1 and G2 phases were counted by the Hoechst intensity (1 for G1 and 2 for G2) in single-cell plot analysis. The thresholds for selecting cells in each cell cycle phase were as follows: G1 (Hoechst < 1.125 and log_2_(EdU) < −3), S (log_2_(EdU) > −3), and G2 (Hoechst > 1.7 and log_2_(EdU) < −3) phases. The number of M phase cells was counted based on the images of Hoechst staining in the same preparations. The G1 phase was subdivided into “low MCM” and “high MCM” states by the value of MCM levels in the early G1 cells as a threshold. To mark cells in the early G1 phase, the “8 h” plot of the pulse-chase experiment were used because the S phase cells pulse-labeled with EdU at 0 h entered the early G1 phase at 8 h chase-incubation (“8 h” in Fig. 2A). The values of log_2_(Mcm3) thresholding “low MCM” and “high MCM” levels were: 0.5 for hTERT-RPE1 (Fig. 2A), 0.3 for HeLa, 0.7 for U2OS, 1.0 for MRC5, and 1.5 for IMR90 (Fig. S3A).

### ChIP assay

ChIP assays were performed as previously described (35,37) with minor modifications. hTERT-RPE1 cells (7.5 × 10^6^ cells) were first immersed in ice-cold 0.2% Triton X-100 containing 20 mM HEPES [pH 7.4], 100 mM NaCl, and 300 mM sucrose for 5 min, and then cross-linked with a 1% formaldehyde solution (Electron Microscopy Sciences, Hatfield, PA, USA) for 10 min at room temperature (∼25°C), followed by the quenching step as previously described (35,37). For immunoprecipitation, mouse monoclonal anti-Mcm3 antibody (M038-3; MBL) and anti-mouse IgG conjugated to Dynabeads M-280 (Thermo Fisher Scientific) were used as the primary and secondary antibodies, respectively.

### ChIP-seq data analysis

The ChIP-seq library of Mcm3 was prepared using the ThruPLEX DNA-seq Kit (Takara Bio), and sequencing was performed using HiSeq1500 (Illumina). The sequenced reads were aligned to the human genome (hg38) using Bowtie software (version 0.12.8; parameter -v3 -m1). To generate input-normalized ChIP-seq signal tracks (bigWig), deepTools software (38) was used (version 3.3.1; bamCompare parameters: -- effectiveGenomeSize 2805636331 --normalizeUsing RPKM --scaleFactorsMethod None -bs 500 --smoothLength 50000). For Orc2, different parameters (bamCoverage: -- effectiveGenomeSize 2701495761 --normalizeUsing RPKM -bs 500 --smoothLength 50000) were used because of a lack of input. The statistics and quality checks of the ChIP-seq data are summarized in Table S1.

### Sucrose gradient fractionation

Fractions of native chromatin were prepared as previously described (39,40) with some modifications. Briefly, hTERT-RPE1 cells (1 × 10^7^ cells) were washed with cold PBS, suspended in buffer A (50 mM HEPES [pH 7.4], 100 mM KCl, 2.5 mM MgCl_2_, 0.1 mM ZnSO_4_, 1 mM ATP, 1 mM DTT, PhosSTOP™ tablets (Roche), and protease inhibitor cocktail (EDTA-free, Nacalai Tesque, Inc.)), and collected by centrifugation (1,300 × *g*, 3 min, 4°C). The cells were then treated with 400 µL of 0.2% Triton X-100 in buffer A, layered on 400 µL of 30% sucrose in buffer A containing 0.2% Triton X-100, and centrifuged at 20,000 × *g* for 10 min at 4°C. The pellet (P1) was collected as a chromatin-bound fraction, and the supernatant (S) was collected as a chromatin-unbound fraction (Fig. 5A). The pellet P1 was washed once with buffer A containing 0.2% Triton X-100 and collected as a pellet by centrifugation (P2). The pellet P2 was resuspended in 300 µL of buffer B (50 mM HEPES [pH 7.4], 10 mM KCl, 2 mM MgCl_2_, 0.1 mM ZnSO_4_, 3 mM ATP, 1 mM DTT, PhosSTOP™ tablets (Roche), protease inhibitor cocktail (EDTA-free, Nacalai Tesque)), and then treated with 200 U/mL Benzonase (Millipore) for 1 h at 4°C to digest DNA and RNA (Fig. 5A). The benzonase-treated P2 fraction (P2+B) was applied to a 5–30% sucrose gradient in buffer A (4.7 mL) and centrifuged at 40,000 rpm (∼128,000 × *g*) for 3 h at 4°C using an MLS-5 rotor (Beckman Coulter). Aldolase (158 kDa) and thyroglobulin (669 kDa) were used as protein size markers for sucrose gradient centrifugation. After centrifugation, the fractions (350 μL each) were collected from the top of the gradient. Proteins in each fraction were concentrated by acetone precipitation, resuspended in buffer A, and subjected to methanol/chloroform precipitation to remove sucrose. The precipitate was dissolved in 2× SDS loading buffer at room temperature (∼25°C) overnight and subjected to Western blot analysis.

### Western blot analysis

Proteins in 2× SDS loading buffer were separated on a 10% SDS polyacrylamide gel and transferred to Immobilon-P PVDF membranes (Merck) using tank blotting (Bio-Rad), with the exception of histone H3 and GAPDH, which were separated on a 15% SDS polyacrylamide gel and transferred using a semi-dry blotting system (ATTO). After blocking with 5% skim milk (Nacalai Tesque), the membranes were incubated with the following primary antibodies: anti-Mcm2 antibody (D7G11; Cell Signaling Technology), anti-GAPDH antibody (14C10; Cell Signaling Technology), anti-histone H3 antibody (1G1, (41)), anti-Cdc6 antibody (sc-9964; Santa Cruz Biotechnology), anti-Cdt1 (D10F11; Cell Signaling Technology), anti-Orc2 antibody (3G6; Santa Cruz Biotechnology), and anti-Cdc45 antibody (D7G6; Cell Signaling Technology). After incubation with peroxidase-conjugated secondary antibodies (GE Healthcare), the signals were detected using ImmunoStar LD (Fujifilm).

## Results

### Single-cell plot analysis to visualize chromatin-bound MCM

To quantify the amount of chromatin-bound MCM in individual human cells, single-cell plot analysis based on fluorescence microscopy was used to measure multiple intracellular components in individual cells. Human cells contain a large soluble pool of MCMs (42-44). Therefore, to quantify chromatin-bound MCM in single-cell plot analysis, we first examined the fixation methods suitable for this analysis. An asynchronous cell population of hTERT-immortalized retinal pigment epithelial (hTERT-RPE1) cells were fixed with two different fixation methods (“direct fixation” or “pre-extraction” in Fig. 1A). Pre-extraction is a method to extract soluble proteins from cells by treating with 0.2% Triton X-100 before fixation, whereas direct fixation is a method to fix the cells without extraction (see Materials and Methods). The fixed cells were fluorescently labeled with Hoechst 33342 (DNA amount indicator), EdU (S phase indicator), and anti-Mcm3 antibody, as described in Materials and Methods. As shown in Fig. 1A, the fluorescence images indicated that Mcm3 staining in cells fixed by the direct fixation method was relatively uniform among the individual cells, whereas that in cells fixed by the pre-extraction method varied. Single-cell plot analysis was performed for the images shown in Fig. 1A (Fig. 1B, see Materials and Methods), and the normalized intensities of EdU (Fig. 1B, left panels) and Mcm3 (Fig. 1B, right panels) in the same cell were plotted against the DNA content determined from Hoechst intensity. EdU-positive cells, representing S phase cells, were plotted as orange dots in the EdU panels, and the same cells were also plotted as orange dots in the Mcm3 panels. In the direct fixation method, the Mcm3 intensities showed relatively small variations ranging from –1.0 to 0.75 on the log_2_ scale with respect to the average intensity throughout the cell cycle (right upper panel in Fig. 1B). In contrast, in the pre-extraction method, the intensities of Mcm3 displayed large variations, ranging from –2.0 to 1.5 on the log_2_ scale during the G1 phase (Fig. 1B). Mcm3 levels were highest at the G1/S transition, decreased during the S phase, and reached the lowest levels in the G2 phase. A similar single-cell plot pattern was obtained for Mcm2 in the single-cell plot analysis (Fig. S1). Such changes in the levels of Mcm3 and Mcm2 during the cell cycle are consistent with previous reports showing the amount of chromatin-bound MCM complex (44-46). Thus, the pre-extraction method is thought to reflect the amount of the MCM complex on chromatin. Based on this result, we used the pre-extraction method for subsequent analyses to detect chromatin-bound MCM complexes.

**Figure 1.**
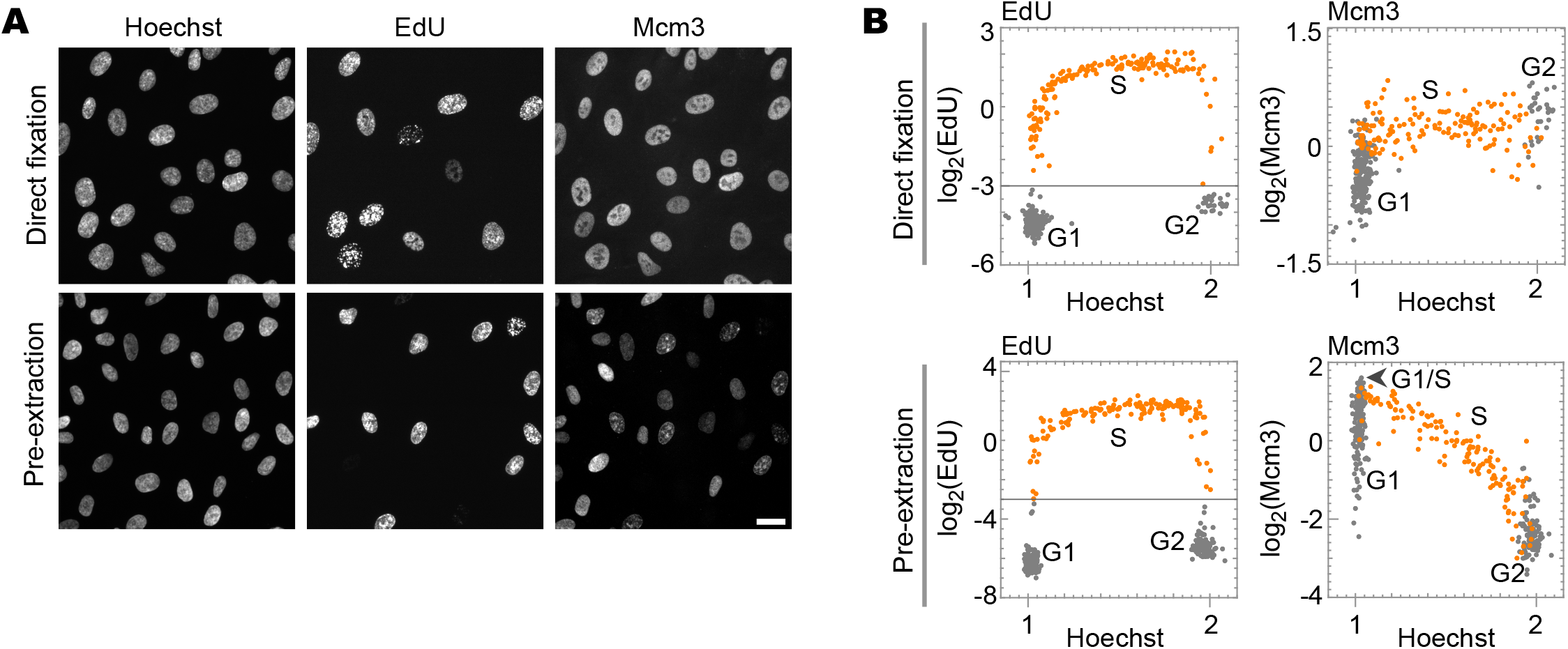
Detection of chromatin-bound MCM by single-cell plot analysis. **A**. Representative fluorescence images of Hoechst 33342, EdU, and Mcm3 in hTERT-RPE1 cells prepared by the direct fixation method (upper panels) or the pre-extraction method (lower panels). Scale bar, 20 µm. **B**. Single-cell plot analysis based on the images in **A**. Each dot represents the intensities of EdU and Mcm3 in a single individual cell plotted against the Hoechst 33342 intensities. The orange dots represent S phase cells based on EdU intensities (log_2_(EdU) > –3). The number of cells examined in each panel is 400.

### Cell-cycle dynamics of MCM revealed by single-cell plot analysis

Using the pre-extraction fixation method, we examined dynamic changes in the levels of chromatin-bound Mcm3 during the cell cycle by pulse-chase experiments. Cells were pulse-labeled with EdU for 30 min, chase-cultured in the medium without EdU for up to 24 h, and then analyzed by single-cell plot analysis (Fig. 2A). Cells undergoing S phase when pulse-labeled with EdU at 0 h (EdU > 0.125; −3.0 on the log_2_ scale) were plotted as orange dots (Fig. 2A, upper leftmost panel, 0 h). In the chase experiments, the orange-marked cells undergoing S phase at 0 h proceeded to different cell cycle stages over time (4–24 h in Fig. 2A). The intensities of Mcm3 in the same cells were plotted against the Hoechst intensity (Fig. 2A, lower panels).

**Figure 2.**
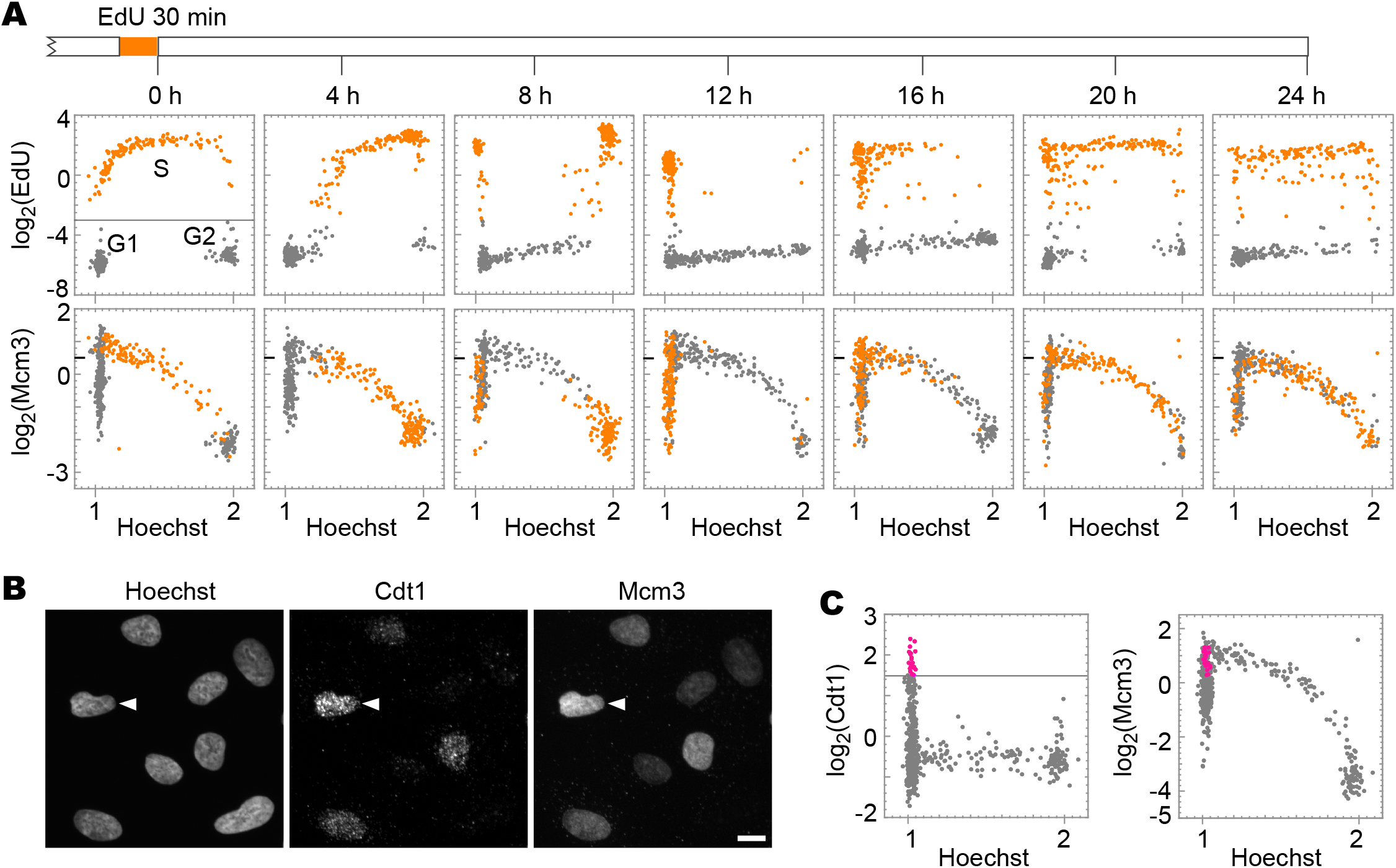
Dynamics of chromatin-bound Mcm3 during the cell cycle. **A**. Single-cell plot analysis of Mcm3 and EdU in hTERT-RPE1 cells in growing phase. The cells pulse-labeled with EdU for 30 min are chase-cultured without EdU. The cells were fixed at the indicated times (timescale on the top) and applied to the single-cell plot analysis for EdU (upper panels) and Mcm3 (lower panels). Each dot represents the intensities of EdU and Mcm3 in a single individual cell plotted against Hoechst 33342 intensities. The orange dots represent cells undergoing S phase at time of 0 h (log_2_(EdU) > −3). The number of cells examined in each panel is 400. **B**. Representative microscopic images of Hoechst, Cdt1, and Mcm3. Arrowheads indicate a cell with high levels of both Cdt1 and Mcm3. Scale bar, 10 µm. **C**. Single-cell plot analysis based on the images in **B**. The pink dots represent G1 phase cells with high Cdt1 levels (log_2_(Cdt1) > 1.5 and Hoechst < 1.125). The number of cells examined in each panel is 400.

In the EdU panels (Fig. 2A, upper panels), 4–8 h after the pulse label, the majority of the orange-marked cells proceeded from the mid/late S phase to G2 phase, as indicated by an increase in the Hoechst intensity. After 8–12 h, a part of the orange-marked cell population proceeded through the M phase and entered the G1 phase, as indicated by low Hoechst intensity, and after 16 h, some cells entered the S phase again. After 20–24 h, most orange-marked cells displayed a middle range of Hoechst intensities with a pattern similar to that of cells at 0 h, consistent with the doubling time in this cell line (approximately 22.5 h) (Fig. 3B). Using the EdU pattern as a cell cycle landmark, we analyzed the cell cycle changes of Mcm3 (Fig. 2A, lower panels). The S phase cells at 0 h were also plotted as orange dots in the Mcm3 panels. Mcm3 levels were the highest (>0.5 on the log_2_ scale) in the early S phase (Hoechst intensity around 1.2) at 0 h and decreased during the late S phase at 4 h. At 8 h, the orange-marked cells were entered into the early G1 phase and showed low to moderate Mcm3 intensity levels (<0.5 on the log_2_ scale; termed as “low MCM” fraction). At 12 h, the Mcm3 intensity levels of orange-marked cells increased to higher levels (>0.5 on the log_2_ scale; termed as “high MCM” fraction), reaching the highest level at 16 h, and then decreased over time. The increase in the Mcm3 levels from the “low MCM” fraction to the “high MCM” fraction suggested that more MCM complexes were recruited to chromatin in the late G1 phase.

**Figure 3.**
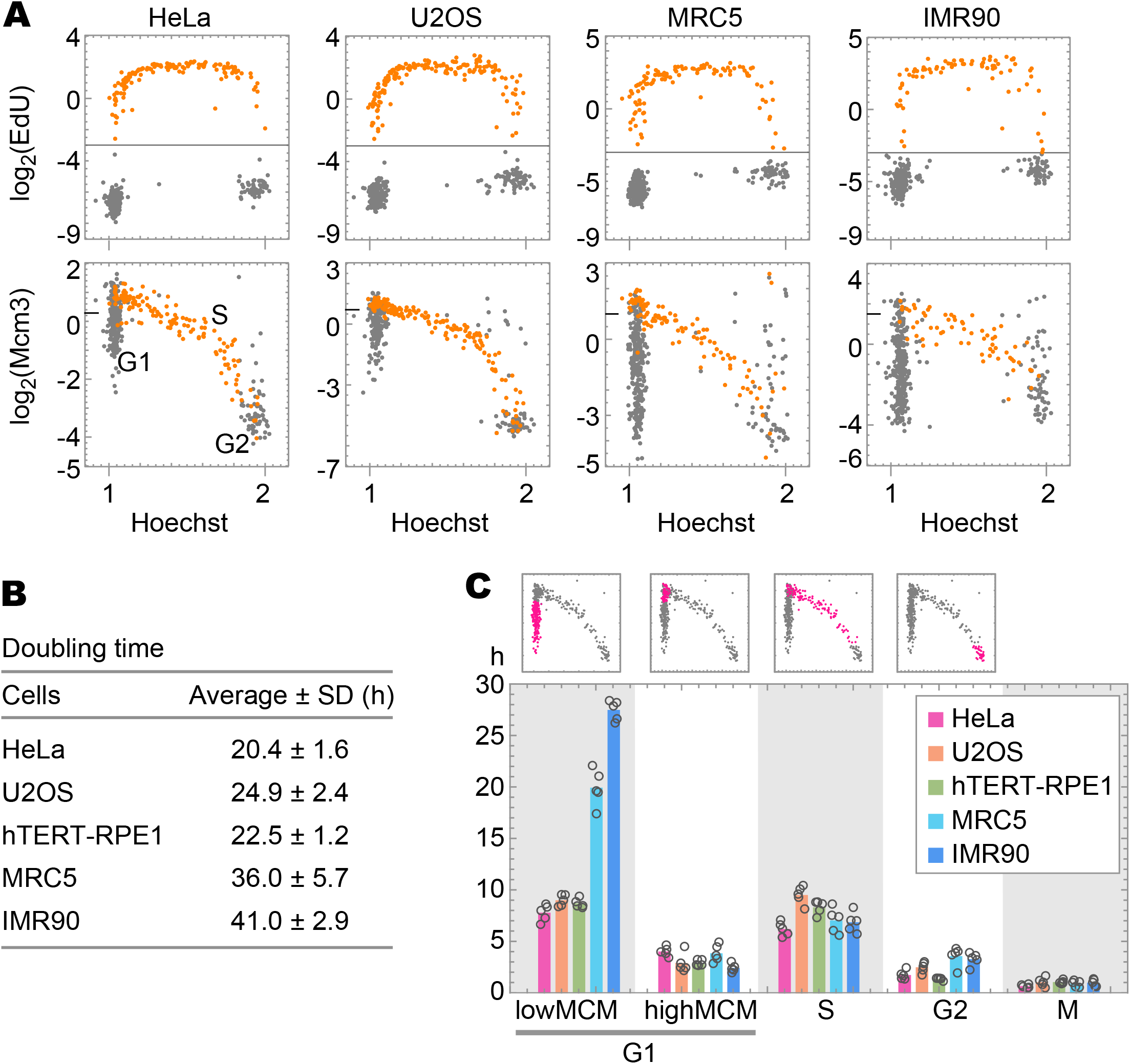
Dynamics of chromatin-bound Mcm3 in various cell lines. **A**. Single-cell plot analysis of Mcm3 and EdU in HeLa, U2OS, MRC5, and IMR90 cells in the growth phase. The cells were pulse-labeled with EdU for 30 min, fixed by a pre-extraction method, and applied to the single-cell plot analysis for EdU (upper panels) and Mcm3 (lower panels). The orange dots represent cells undergoing S phase at 0 h (log_2_(EdU) > −3). The number of cells examined in each panel is 450. The tick mark on the vertical axis of Mcm3 indicates a border between low and high levels. **B**. Doubling times of HeLa, U2OS, hTERT-RPE1, MRC5, and IMR90 cells in the growth phase. The doubling time was determined from growth curves in five independent experiments. **C**. Duration for each phase of the cell cycle: G1 (low MCM, high MCM), S, G2, and M phases. Graphs on the top are examples of single-cell plot analysis in hTERT-RPE1 cells; magenta dots represent cells assigned for each of the phases indicated at the bottom. An estimation of the duration of each cell cycle phase is described in Materials and Methods. The averages are plotted as bars and each data point (n = 5) are shown as open circles.

To confirm “high MCM” cells in the late G1 phase, we examined the profile of phosphorylated Mcm2 at the 108^th^ serine residue (Mcm2-S108ph) in relation to those of Mcm3 and EdU (Fig. S2A, B). The cells with a “high Mcm3” signal (arrows in Fig. S2A and pink dots in Fig. S2B) showed relatively higher levels of Mcm2-S108ph but showed no EdU signals. The phosphorylation of Mcm2 occurs immediately before and during the S phase (47,48). Thus, this result suggests that “high Mcm3” cells are indeed in the late G1 phase.

To understand whether the high Mcm3 levels in the late G1 phase reflect loading to chromatin, we examined the profile of Cdt1 protein in relation to that of Mcm3 (Fig. 2B, C and Fig. S2C), since Cdt1 is known to promote the loading of MCM complexes onto chromatin (17). Single-cell plot analysis of Cdt1 and EdU revealed that the Cdt1 levels varied from low to high in G1 cells (Hoechst < 1.125) (gray and pink dots in Fig. 2C left panel) but remained low in S phase cells (orange dots in Fig. S2C). The single-cell plot analysis of Cdt1 and Mcm3 revealed that the cells with the highest levels of Cdt1 (>1.5 on the log_2_ scale; pink dots in Fig. 2C left panel) were plotted to the “high Mcm3” fraction (>0.5 on the log_2_ scale; pink dots in Fig. 2C right panel). This result suggests that Mcm3 proteins are more loaded onto chromatin in these high-Cdt1 cells.

### Cell-cycle dynamics of MCM in various cell types

We also examined the cell-cycle dynamics of chromatin-bound MCM for various cell types using pulse-chase experiments. Two cancer cell lines, HeLa and U2OS, and two normal cell lines, MRC5 and IMR90, were evaluated. Cells were pulse-labeled with EdU for 30 min (Fig. 3A) and cultured in the medium without EdU for 8 h (Fig. S3A). The MCM levels in “8 h” cells were used to determine the values thresholding between the “low MCM” and “high MCM” states (see Materials and Methods for details). EdU-positive cells (EdU > −3.0 on the log_2_ scale) undergoing S phase at 0 h were plotted as orange dots against the Hoechst intensity (Fig. 3A, upper panels). The intensities of Mcm3 in the same cells were also plotted against Hoechst intensity (Fig. 3A, lower panels). The Mcm3 levels varied from low to high in the G1 phase, highest at the G1/S transition, decreased during the S phase, and reached lower levels in the G2 phase in all the cell lines examined. The plot patterns from these cell lines were similar to those of hTERT-RPE1 cells, suggesting that cell cycle dynamics at MCM levels may be common in human cells. However, there were some differences among the cell lines; that is, cell population with the “low MCM” state was small in hTERT-RPE1 (36%), HeLa (38%), and U2OS cells (36%), whereas it was large in MRC (53%) and IMR90 (67%) (gray dots at the position of G1 at 0 h, Fig. 3A lower panels). As the MRC5 and IMR90 cells showed longer doubling times (Fig. 3B), we investigated which cell cycle stages were prolonged in these cells (Fig. 3C). The duration of each cell cycle stage (“low MCM”-G1, “high MCM”-G1, S, G2, and M) was determined (Fig. 3C), as described in Materials and Methods. Although the duration of “high MCM” G1 phase was 2–4 h in all the cell lines examined, the duration of the “low MCM” G1 phase was 20–27 h in MRC5 and IMR90 cells and 8–9 h in hTERT-RPE1, HeLa, and U2OS cells (Fig. 3C). Therefore, the prolonged doubling time was attributed to the high proportion of “low MCM” in the G1 phase. These results indicate that the normal cells have a prolonged G1 phase in a “low MCM” state.

To understand the prolonged G1 phase with “low MCM” levels in these normal cells, we examined the levels of phosphorylated retinoblastoma protein (Rb) at Ser807/811, which is necessary for cell cycle progression during the G1 phase (49-51). A large population of the G1 phase of MRC5 had low levels of phosphorylated Rb (gray dots in the square region in the right panel of Fig. S3B). In these cells, the levels of phosphorylated Rb were lower than those in the newly entering G1 cells (orange dots in Fig. S3B), which were determined by EdU signals at 8 h after pulse labeling (see Materials and Methods for details). Cells with low levels of phosphorylated Rb were rarely observed in proliferating hTERT-RPE1 cells (square region in the left panel of Fig. S3B). These results suggest that MRC5 cells in the G1 phase have an additional pausing phase during cell cycle progression.

### MCM dynamics under G1 arrest

To investigate the MCM states in G1 arrested cells, cell cycle progression was halted under two conditions in hTERT-RPE1 cells (Fig. 4A, B). One condition involved contact inhibition by culturing the cells to confluence. The other condition was via depletion of Set8, the methyltransferase for histone H4K20 monomethylation (H4K20me1) (52,53) by siRNA treatment. We used a relatively long siRNA treatment time of more than 40 h because it was required to arrest the cell cycle (Fig. S4) (see Supplementary Results for shorter Set8-siRNA treatment). In both confluent and Set8-siRNA conditions, the number of EdU-positive cells was low and most of the cells showed low Hoechst intensities (Fig. 4B), suggesting that these cells were arrested in the G1 phase. Remarkably, the cells treated with Set8-siRNA showed a complete decrease in the levels of histone H4K20me1 (<–3 on the log_2_ scale with respect to the average intensity of control cells), whereas the confluent cells showed low to moderate levels of H4K20me1 from −3 to 0 (on the log_2_ scale). In this regard, the Set8-siRNA-treated cells showed high MCM levels (>–0.5 on the log_2_ scale) that were sufficient for entry into the S phase, whereas the Mcm2 intensity was low to moderate from −4.5 to – 0.5 (on the log_2_ scale) in the confluent cells. These results indicate that confluent hTERT-RPE1 cells were arrested at a “low MCM” level similar to the normal cell lines (MRC5 and IMR90), and Set8-siRNA treated cells were arrested at a “high MCM” level in the G1 phase. In Set8-siRNA treated cells, the levels of H4K20me2/me3 were comparable to those of control siRNA (single-cell plot analysis in Fig. S4B). This result suggests that pre-existing H4K20me1 was converted to H4K20me2/me3, leading to a high MCM state in these cells.

**Figure 4.**
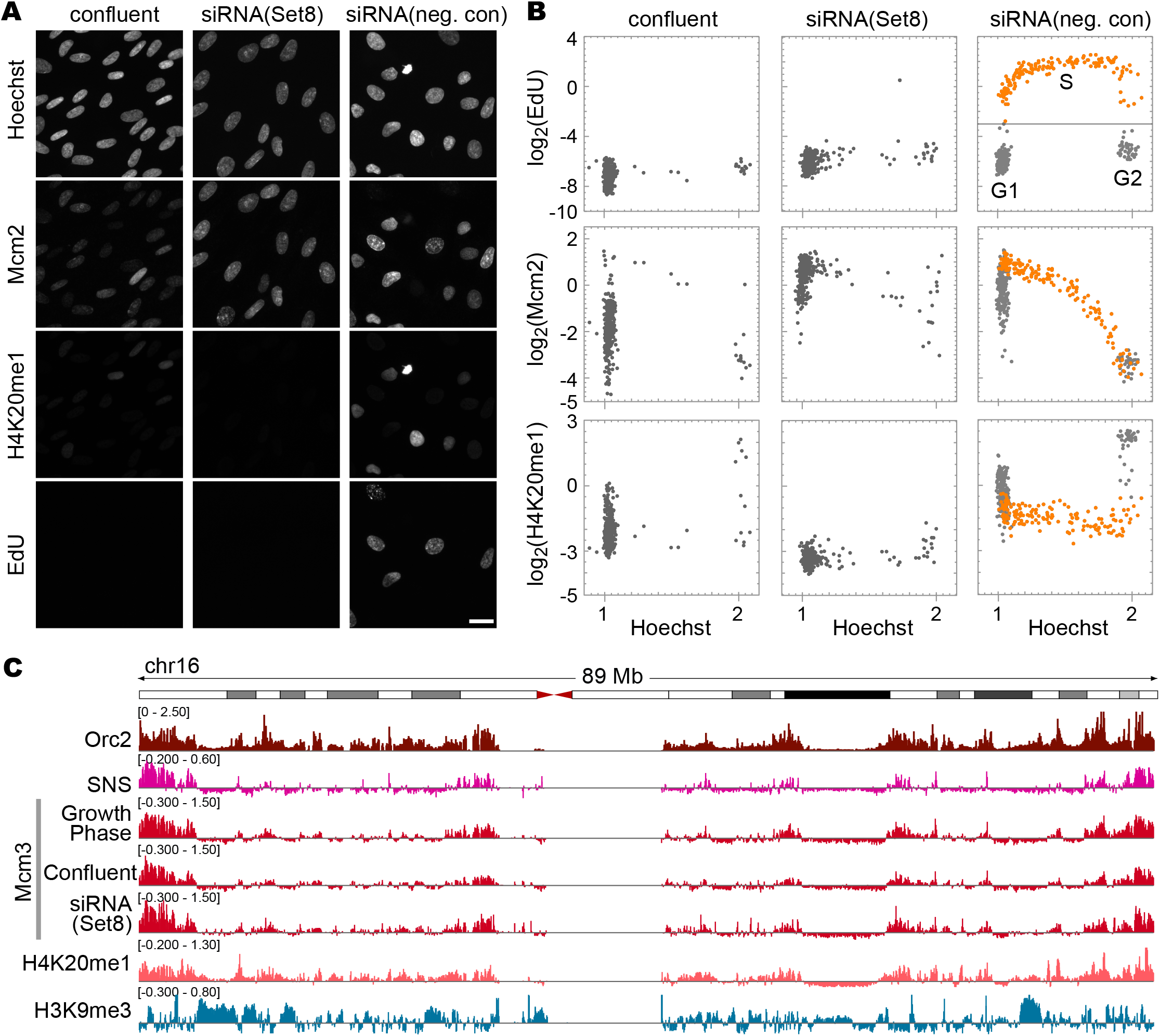
MCM dynamics under conditions of G1 arrest in hTERT-RPE1 cells. **A**. Representative microscopic images of Hoechst, Mcm2, histone H4K20me1, and EdU in hTERT-RPE1 cells under confluent conditions or treated with either Set8-siRNA or negative control-siRNA. Scale bar, 20 µm. **B**. Single-cell plot analysis based on the images in **A**. Each dot represents the intensities of Mcm2, histone H4K20me1, and EdU in a single individual cell plotted against Hoechst 33342 intensities. The orange dots represent S phase cells based on EdU intensities (log_2_(EdU) > −3) in siRNA (negative control). The number of cells examined in each panel is 400. **C**. Genome-wide analysis of Mcm3 localization characterized using ChIP-seq. An example of chromosome 16: enrichment of Mcm3 (growth phase, confluent, and Set8-siRNA) obtained in this study is shown together with that of Orc2 (K562 cells; GSM1717888), SNS (IMR-90 cells; GSM927235), and H4K20me1 (HeLa-S3 cells; GSM733689), and H3K9me3 (IMR90 cells; GSM1528890) obtained from public database. Vertical axes are shown on the linear scale for Orc2 (RPKM_ChIP) and on the log_2_ scale for the others (RPKM_ChIP/RPKM_Input).

### Genome-wide distribution of MCM3 proteins

To examine the differences in MCM binding preferences in the genome during early or late G1 phases (corresponding to the “low” and “high” MCM state, respectively), we first determined the genomic distribution of MCM binding using chromatin immunoprecipitation followed by DNA sequencing (ChIP-seq) analysis with the Mcm3 antibody. The asynchronous population of cells in the growth phase was pre-extracted with Triton X-100 before fixation, followed by normal preparation for ChIP-seq experiments (35,37). Chromatin prepared from 7.5 × 10^6^ cells produced significant enrichment of the Mcm3 peaks (Fig. 4C, Table S1), which was similar to the Mcm7 peaks (54). Then, the Mcm3 profile was compared with previous reports on Orc2 ChIP-seq data (55) and SNS-seq data (sequences of short, nascent DNA single strands reflecting replication initiation) (56). Orc2 is required for the loading of MCM complexes and is regarded as a marker of replication origin (16). The pattern of the Mcm3 peaks was similar to that of the Orc2 and SNS peaks (Fig. 4C and Fig. S5A), indicating that the MCM complex accumulated at the replication origin. However, in some chromosomes that showed late replication initiation (57), SNS and Mcm3 accumulated less, whereas the Orc2 peaks were detectable, for example, in chromosome 6 (Fig. S5B). Because Orc2 remains on the chromatin throughout the cell cycle (58,59), these results suggest that MCM abundance is a marker of early replication rather than Orc2-marked replication origin.

We also examined the Mcm3 distribution in G1-arrested cells (confluent cells and Set8-siRNA-treated cells) using ChIP-seq experiments. The genome-wide localization patterns of Mcm3 were almost indistinguishable among these cells (Fig. 4C, Table S1), indicating that variations shown in immunostaining do not influence the genomic localization pattern of MCM.

### Single and double MCM hexamers on chromatin

The increase in the amount of MCM during the G1 phase revealed by the immunostaining experiment may reflect the conversion of a single to double hexamer of the MCM complex. To evaluate this possibility, we examined the size of the chromatin-bound MCM complex using a sucrose gradient centrifugation method. Chromatin-bound fractions were prepared from hTERT-RPE1 cells according to a previously described method (39,40) with some modifications (Fig. 5A) (see Materials and Methods for details). First, the cells were treated with Triton X-100 to extract soluble proteins and centrifuged through a sucrose cushion to separate the pellet fraction (P1) and supernatant fraction (S). The P1 and S fractions were evaluated by Western blot analysis (Fig. 5B). We examined cells grown under different conditions: growing cells, growth-arrested cells by confluent culture, and Set8-depleted cells by siRNA treatment. In all conditions tested, the S fraction contained chromatin-unbound proteins, such as GAPDH, but not chromatin-bound proteins, such as histone H3; on the other hand, the P1 fraction contained histone H3 but not GAPDH (Fig. 5B, middle and bottom panels). Approximately the same amount of Mcm2 as in the P1 fraction was detected in the S fraction from growing and confluent cells, although faint Mcm2 signals were detected in the S fraction from Set8-depleted cells (Fig. 5B, top panel). These results indicate that the chromatin-bound fraction (P1) was effectively separated from the chromatin-unbound fraction (S) by this fractionation method. Then, the chromatin-bound P1 fraction was further washed with Triton X-100 and collected as a pellet (P2). The P2 fraction was treated with Benzonase to digest DNA and RNA (Fig. 5A). The reaction mixture (P2+B) was subjected to sucrose gradient centrifugation to assess the size of the MCM on the chromatin.

**Figure 5.**
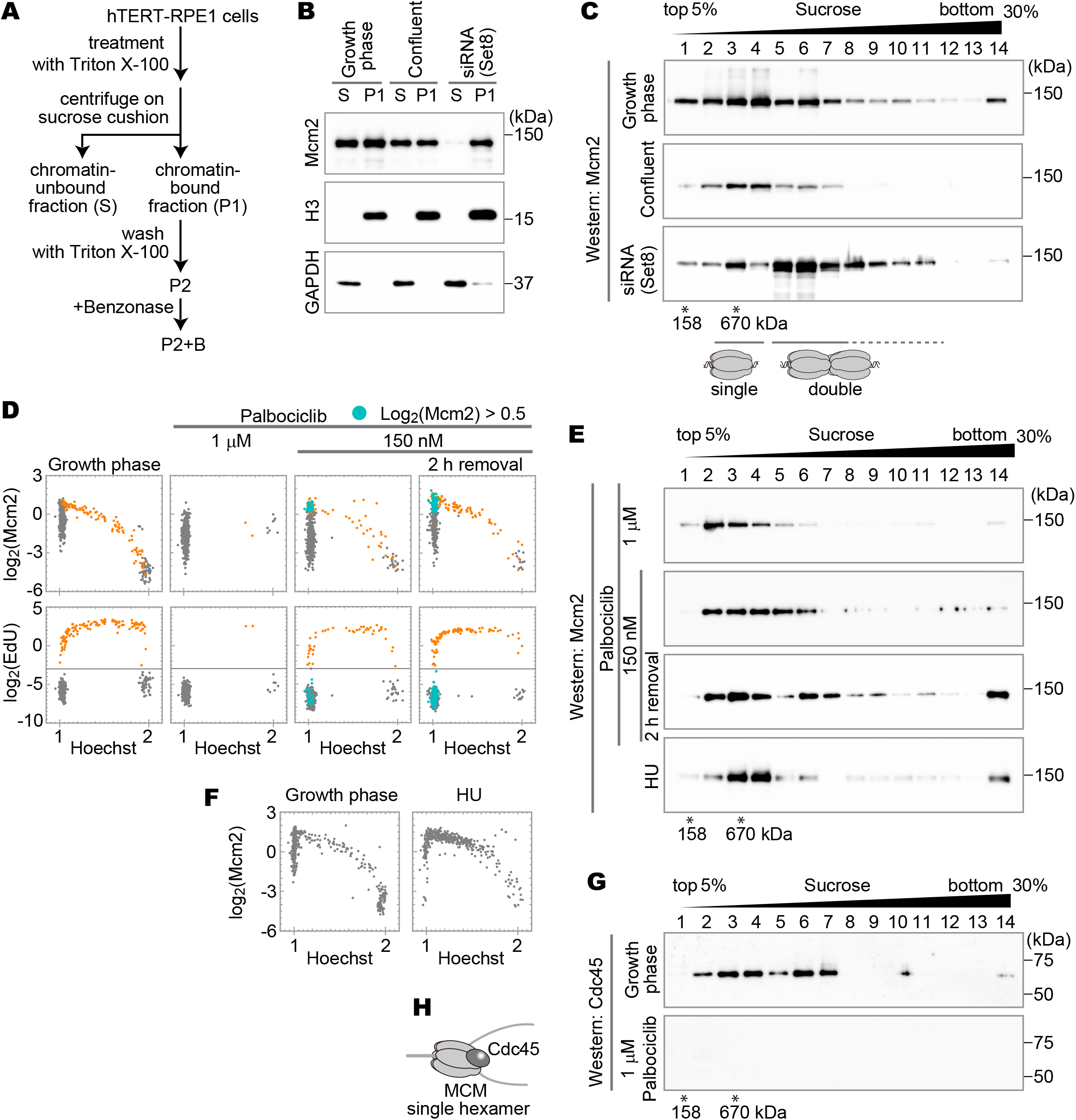
Identification of MCM hexamer states. **A**. A schematic diagram for preparation of chromatin-bound fractions. Chromatin-bound fractions (P1, P2 and P2+B) and chromatin-unbound fraction (S) were prepared as indicated. **B**. Western blot analysis of P1 and S fractions using antibodies against Mcm2, histone H3, and GAPDH. The fractions prepared from the growth phase cells, confluent cells, and the cells treated with Set8-siRNA were applied. **C**. Western blot analysis of sucrose gradient fractions using anti-Mcm2 antibody. The P2+B fractions prepared from the growth phase cells, confluent cells, and Set8-siRNA-treated cells were fractionated through a linear 5–30% sucrose gradient. Molecular sizes derived from the sucrose gradient are indicated below the blots. The result of the S fraction is shown in Fig. S6A. **D**. Single-cell plot analysis of Mcm2 (upper panels) and EdU (lower panels) in the growth phase cells, palbociclib-treated cells (1 μM or 150 nM), and 2 h after the release from the 150 nM palbociclib arrest as indicated. The orange dots represent S phase cells based on EdU intensities (log_2_(EdU) > −3). The blue dots represent G1 cells with high MCM levels (log_2_(Mcm2) > 0.5, Hoechst < 1.125 and (log_2_(EdU) < −3). Longer times (up to 8 h) after the release from the 150 nM palbociclib arrest are also shown in Fig. S6D. **E**. Western blot analysis of sucrose gradient fractions using anti-Mcm2 antibody. The P2+B fractions were prepared from palbociclib-treated cells (1 μM or 150 nM), 2 h after the release from the 150 nM palbociclib arrest, and HU-treated cells as indicated, and fractionated through a linear 5–30% sucrose gradient. **F**. Single-cell plot analysis of Mcm2 in HU-treated cells compared with the growth phase cells. The number of cells examined in each panel is 450. **G**. Western blot analysis of sucrose gradient fractions using anti-Cdc45 antibody. The P2+B fractions prepared from the growth phase cells and 1 μM palbociclib-treated cells were fractionated through a linear 5–30% sucrose gradient. **H**. Cartoon of a single MCM hexamer with Cdc45.

Sucrose gradient centrifugation showed that chromatin-bound Mcm2 was detected in fraction 6 in the growing cells, in addition to a peak in fractions 3 and 4 (Fig. 5C, upper panel). On the other hand, in the confluent cells, the majority of chromatin-bound Mcm2 was found in fractions 3–4, although weak signals appeared in the larger fractions (up to fraction 7) (Fig. 5C, middle panel). In the Set8-siRNA treated cells (Fig. 5C, bottom panel), the Mcm2 proteins were abundantly present in fractions 5–6. From the predicted molecular weight of the single MCM hexamer (∼545 kDa), it is likely that a smaller peak (fraction 3) corresponds to a single MCM hexamer and that the larger peak (fraction 6) corresponds to a double MCM hexamer. These results are consistent with the results of single-cell plot analysis showing low and high levels of Mcm2 in the confluent and Set8-siRNA treated cells, respectively (Fig. 4B).

### Single and double MCM hexamers in G1 arrest cells

To further examine the MCM hexamer states during the progression through the G1 to S phase, the cells were synchronized at the G1 phase. To this end, palbociclib was used, since this reagent is known to arrest the cells at the G1 phase by inhibiting Cdk4/6 activity (60), which is required for cell cycle progression from G1 to S phase. Synchronized cells were evaluated by single-cell plot analysis (Fig. 5D) and sucrose gradient fractionation (Fig. 5E). Cells treated with 1 µM palbociclib were arrested at the G1 phase and exhibited the “low MCM” state in single-cell plot analysis (Fig. 5D). In the sucrose gradient analysis, the chromatin-bound Mcm2 was fractionated in fractions 2–3 (Fig. 5E), slightly smaller than those detected in the growth phase. In these cells, the expression levels of Cdc6 and Cdt1 (chromatin loading factors of MCM) were greatly reduced (Fig. S6B), explaining the “low MCM” state under this condition. When cells were treated with a lower concentration (150 nM) of palbociclib, the cells were mildly arrested at the G1 phase with “low” and “high MCM” states (Fig. 5D, high Mcm2 cells plotted as blue dots). Under these conditions, the Cdc6 and Cdt1 proteins were expressed at levels similar to those observed in growth phase cells (Fig. S6B). Sucrose gradient analysis showed that the Mcm2 signal in the arrested cells was detected in fractions 2–6 (Fig. 5E, second row; see Fig. S6C for the sample evaluation). After the removal of palbociclib, these cells progressed from the G1 phase into the S phase (Fig. 5D and S6D, orange dots of EdU-positive). The number of cells with the “high MCM” state increased from 7% of the total cells to 30% for 2 h after the removal (Fig. 5D, blue dots in the upper right panel). Sucrose gradient analysis showed that the Mcm2 signal was detected in the larger fractions (fractions 6–7, double MCM hexamer size) and fractions 2–4 (single hexamer size) at 2 h after the removal of palbociclib (Fig. 5E, third row). We also examined the S phase-arrested cells to understand MCM states during the S phase. Hydroxyurea (HU) treatment was used to arrest cells in the S phase. Single-cell plot analysis showed that HU-treated cells were effectively arrested in the early S phase (Fig. 5F). Sucrose gradient analysis showed that the Mcm2 signals appeared in fractions 3–4 (a single MCM hexamer size), but not in fraction 6–7 (a double MCM hexamer size) (Fig. 5E, bottom row). These results suggest that the transition from single to double MCM hexamer status occurs before the S phase or late G1 phase.

The larger MCM complexes detected in the sucrose gradient may reflect the association of additional DNA replication initiation factors. To evaluate the state of the MCM complex in sucrose gradient fractions, we examined Cdc45. Cdc45 binds to double MCM hexamers at the replication origin to initiate DNA replication (61-63). Indeed, Cdc45 was detected mostly in the S phase by single-cell plot analysis (Fig. S7A). In the sucrose gradient fractions from growing cells, Cdc45 was detected in both fractions 3–4 and 6–7 containing smaller and larger MCM complexes, respectively (Fig. 5G, upper panel). This result suggests that the larger MCM complex detected by the Mcm2 antibody in the sucrose gradient fractionation (Fig. 5C and E) is not a result of the association of additional factors with the smaller MCM complex. In contrast, Cdc45 was not detected in the fractions from 1 µM palbociclib-arrested G1 cells (Fig. 5G, lower panel) and from the G1-arrested cells in confluent culture (Fig. S7B, C), suggesting that the Cdc45-positive fractions reflect S phase MCM complexes. Because Cdc45-bound double MCM hexamers are converted to single MCM hexamers as DNA replication progresses, Cdc45-positive single MCM hexamers likely reflect a single MCM complex existing at the replication fork progressing during DNA replication (Fig. 5H).

### MCM hexamer states and levels of histone H4K20 methylation

To elucidate the relationship between the levels of MCM and histone H4K20 methylation in hTERT-RPE1 cells, asynchronized cells were co-immunostained with antibodies against Mcm2, H4K20me1, and H4K20me2/me3 (Fig. 6A and Fig. S8A) and subjected to single-cell plot analysis (Fig. 6B and Fig. S8B). As shown in the top row of Fig. 6B, the G1 phase cells with high levels of histone H4K20me1 (>0.3 on the log_2_ scale) were selected and plotted as light blue dots (left), and the same cells were also plotted as light blue dots in the H4K20me2 (center) and Mcm2 (right) panels, showing relatively low levels of H4K20me2 and Mcm2 among the G1 cells. As shown in the middle row of Fig. 6B, cells with low H4K20me2 levels (<–0.1, on the log_2_ scale; blue dots) were selected (center); these cells showed high H4K20me1 levels (left) and low Mcm2 levels (right). In the bottom row of Fig. 6B, G1 cells with high Mcm2 levels (>0.5 on the log_2_ scale; pink dots) were selected (right); these cells showed low H4K20me1 levels (left) and high H4K20me2 levels (center). A scatterplot of these dots against H4K20me1 and Mcm2 intensities indicated that the light blue (high H4K20me1 levels) and pink (high Mcm2 levels) dots were mutually exclusive (Fig. 6C, top). In addition, the blue (low H4K20me2 levels) and pink (high Mcm2 levels) dots were mutually exclusive (Fig. 6C bottom). Similar results were also obtained for H4K20me3 (Fig. S8B, C), indicating that the decrease in H4K20me1 levels reflects an increase in H4K20me2/me3. These results indicated that when the H4K20me1 level was high, the MCM on chromatin was in a single hexamer state, and when the MCM was in a double hexamer state, the H4K20me2/me3 level was high (Fig. 6C and Fig. S8C).

**Figure 6.**
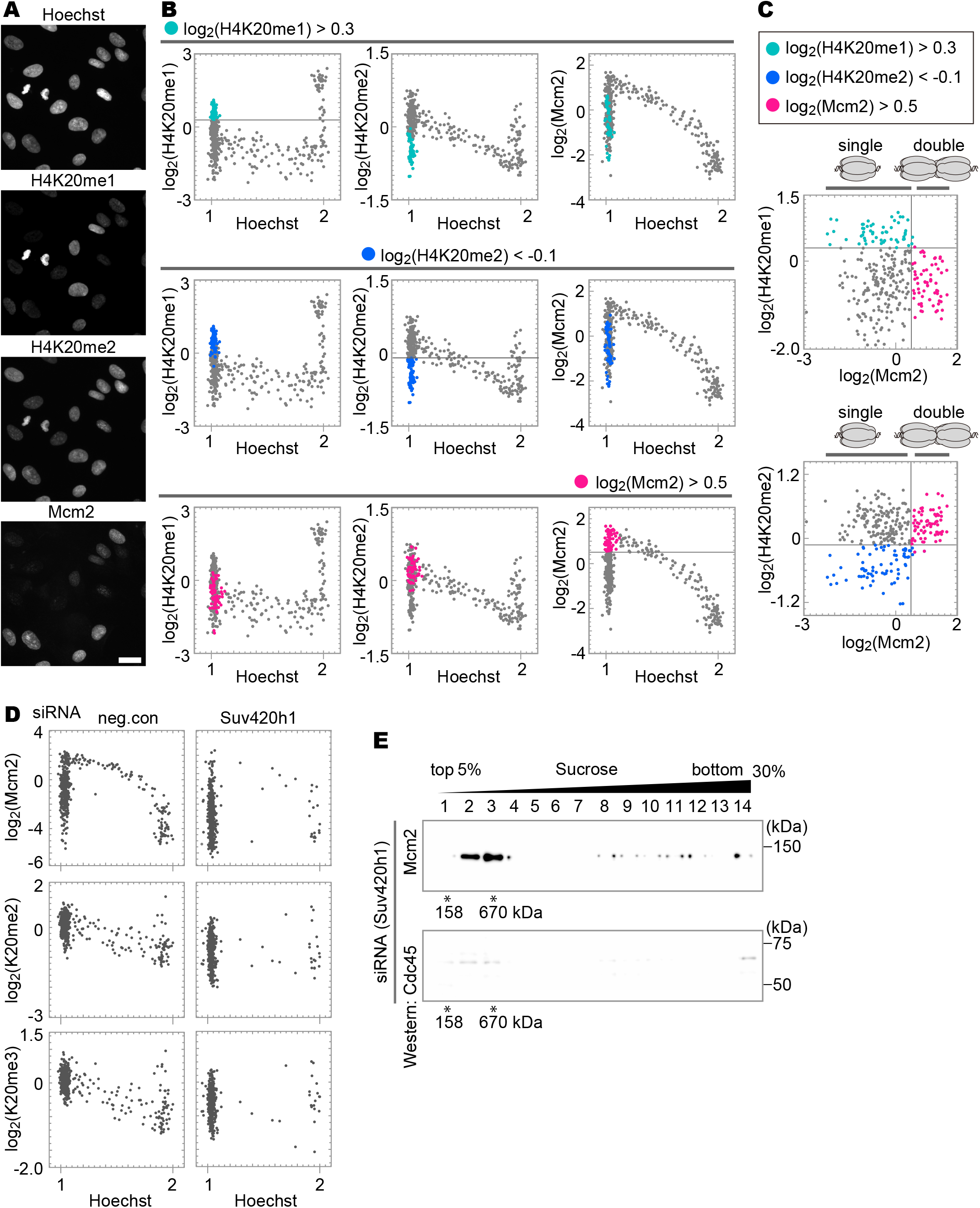
Relationship between histone H4K20 methylation levels and MCM hexamer states. **A**. Representative microscopic images of Hoechst, H4K20me1, H4K20me2, and Mcm2. Scale bar, 20 µm. **B**. Single-cell plot analysis of H4K20me1, H4K20me2, and Mcm2. The G1 cells with high levels of H4K20me1 (Hoechst < 1.125 and log_2_(H4K20me1) > 0.3) are shown in light blue in the upper panel, those with low levels of H4K20me2 (Hoechst < 1.125 and log_2_(H4K20me2) < –0.1) in blue in the middle panel, and those with high levels of Mcm2 (Hoechst < 1.125 and log_2_(Mcm2) > 0.5) in pink in the lower panel. **C**. Scatterplots between H4K20me1 and Mcm2 intensities, and between H4K20me2 and Mcm2 intensities based on **B. D**. Single-cell plot analysis of Mcm2, H4K20me2, and H4K20me3 in cells treated with either control-siRNA, or Suv420h1-siRNA. The number of cells examined in each panel is 450. **E**. Western blot analysis using anti-Mcm2 (upper) and anti-Cdc45 (lower) antibodies following fractionation with a linear 5–30% sucrose gradient. The cells treated with Suv420h1-siRNA were used.

To confirm that the conversion of H4K20me1 to H4K20me2/me3 is necessary for the transition from the single to the double state of MCM hexamers, the cells were treated with siRNA of Suv420h1, the enzyme responsible for this conversion. Compared with the control cells, the Suv420h1-siRNA-treated cells showed lower H4K20me2/me3 levels (Fig. 6D, right middle and lower panels) and a “low MCM” state (Fig. 6D, right top panel). We examined the states of the MCM complex by sucrose gradient ultracentrifugation for the chromatin-bound P2+B fraction from the Suv420h1-siRNA-treated cells (Fig. 6E). The MCM proteins were detected in fractions 2–3, suggesting only a single MCM hexamer. These fractions contained no detectable amount of Cdc45, indicating a single MCM hexamer in the early G1 phase. These results suggest that the conversion of H4K20me1 to H4K20me2/me3 is essential for pre-RC formation in hTERT-RPE1 cells.

## Discussion

In this study, we demonstrated that MCM complexes on chromatin convert from a single hexamer to a double hexamer in association with the di-/tri-methylation of histone H4K20 toward S phase entry. In combination with previous reports (as reviewed in (64)), our results propose the following model (Fig. 7): Mcm2-7 association and loading occur during the G1 phase. The MCM complex forms a single hexamer without Cdc45 in the early G1 phase (state 1). These complexes are subsequently transformed into double hexamers before entering the S phase (state 2). Single-to-double hexamer transformation is inhibited in the cells arrested by treatment with 1 μM palbociclib. Then, Cdc45 is loaded onto the MCM double hexamer to initiate DNA replication in the S phase (state 3). As Set8 is required for S phase progression (25,26), Set8-depleted cells are arrested around the G1/S transition, retaining the Mcm2-7 double-hexamer state. Once DNA replication is initiated, the MCM double hexamer containing Cdc45 converts to the MCM single hexamer containing Cdc45 (state 4) as the replication forks progress as in growth phase cells. The transition of the single to double hexamer of MCM is triggered by the conversion of H4K20me1 to H4K20me2/me3.

**Figure 7.**
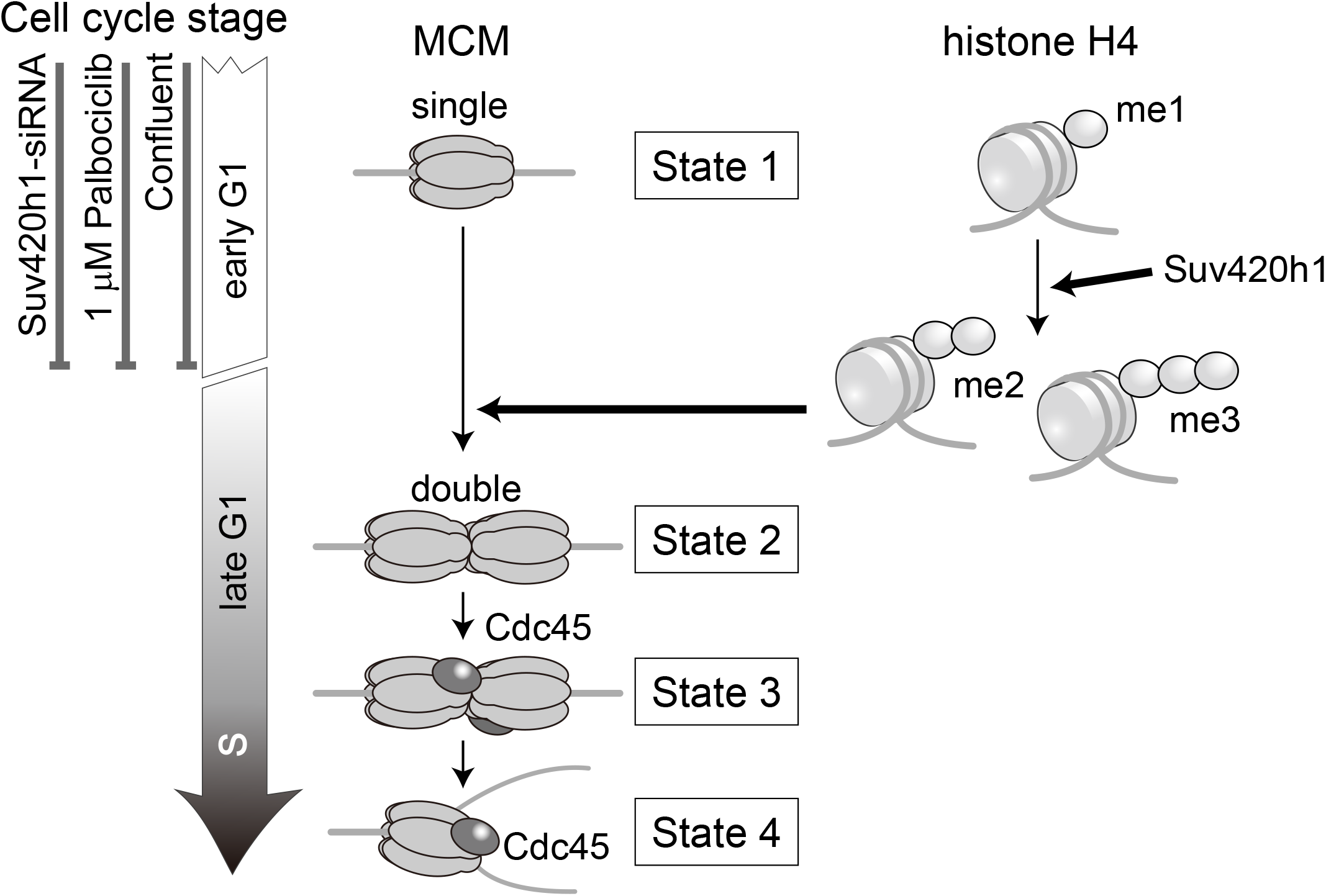
A state model of the MCM hexamer during G1/S phases. The conversion of histone H4 methylation from mono to di-/tri triggers the transition from the single to double state of MCM hexamers.

Normal cells (MRC5 and IMR90) or hTERT-RPE1 cells arrested via contact inhibition showed that cell populations in the single MCM hexamer state increased, compared with HeLa and U2OS cancer cells, indicating that normal cells pause at the single MCM hexamer state longer than cancer cells during the G1 phase. In comparison, the duration of the double hexamer state in the cell cycle was unchanged across cell types. These results suggest that MCM on chromatin pauses at the single hexamer state in the G1 phase and can stay halted if conditions favor a non-proliferating state. In contrast, once the double hexamer forms, the cell cycle proceeds to enter the S phase for cell proliferation. Therefore, halting the single MCM hexamer state is a limiting step in the decision of cell proliferation or quiescence. The transition from the single to double hexamer state of MCM can be affected by the activation of several factors associated with G1 arrest (2,10,65). In fact, when the cells were treated with a higher concentration of palbociclib, the MCM complex remained in the single hexamer state without conversion to the double hexamer. Palbociclib treatment inhibits the activity of Cdk4/6, leading to the inactivation of the Rb protein (60), suggesting that the conversion of the MCM state from a single to double hexamer is regulated by G1 progression. Rb is mutated or functionally inactivated in the majority of cancer cells (66,67), supporting our results, which show that cancer cells pass through the single-to-double MCM hexamer conversion more quickly than normal cells.

Histone modifications, especially methylation at histone H4K20, may be an important factor in regulating the loading of the second MCM complex. Our results indicate that the levels of histone di-/tri-methylation at H4K20 increased prior to the loading of the second MCM hexamer (Fig. 6 and Fig. S8), suggesting that the conversion of H4K20me1 to H4K20me2/me3 is required for the loading of the second MCM hexamers. As the loading of the second MCM hexamer is required for G1 progression to the S phase, the conversion of H4K20me1 to H4K20me2/me3 seems to be an important factor in determining S phase progression. Although the factors that cause the transition of the histone modification state remain unknown, our findings elucidate the relationship between the process of pre-RC formation and histone modifications during the G1 phase, and therefore provide new insights into the mechanism of the proliferation-quiescence decision.

## Supporting information

Supplementary Results, Figure S1-S8 & Table S1

## Data availability

ChIP-seq data were submitted to Gene Expression Omnibus (GEO) with the accession number: GSE157839. Further data are available from the corresponding authors upon reasonable request.

## Acknowledgements

We also thank Drs. Keiichi Namba and Tomoko Miyata (Osaka University) for their kind supports in protein biochemistry experiments.

