## Supplementary Results, Figure S1-S8 & Table S1 for "Chromatin loading of MCM hexamers is associated with di-/tri-methylation of histone H4K20 toward S phase entry"

**Supplementary Figure S1-S8**

**Supplementary Table S1**

### Supplementary Results

#### Effect of Set8-siRNA on MCM hexamer states in shorter treatment times

In the experiments that arrested cells at the G1 phase, the cells were treated with Set8-siRNA for more than 40 h (Fig. 4B, 5C). However, this relatively long treatment with Set8-siRNA may cause an abnormal G1 phase, as previously reported (24). To avoid this, a shorter Set8-siRNA treatment (18 h up to 26 h) was used to evaluate the effect of Set8 depletion on the MCM hexamer states. H4K20me1 levels were effectively reduced in cells treated with Set8-siRNA for 18 h, although not completely lost (Fig. S4A, 0 h, bottom row). Under these conditions, the cell cycle of some cells was not arrested (Fig. S4A, 0 h). To monitor the cell cycle progression of the cells treated with Set8-siRNA, we conducted a pulse-chase experiment in the presence of Set8-siRNA. In the cells treated with Set8-siRNA for another 8 h (26 h in total), the H4K20me1 levels decreased to lower levels than those in the cells treated for 18 h, although they remained higher than those in the cells treated for more than 40 h. Additionally, cells undergoing S phase at 0 h (EdU pulse-labeled cells at 0 h) were in the early G1 phase (blue-marked, Hoechst $< 1.125$, EdU $> 0$ on the $\log_2$ scale; the squares in the EdU panels in Fig. S4A). In these blue-marked Set8-siRNA-treated cells, the Mcm2 levels remained within a range from $-1$ to $0.5$ (on the $\log_2$ scale; blue dots in the Mcm2 panels in Fig. S4A), suggesting that the MCM complex still remains in a single hexamer state. These results are consistent with our finding that the MCM complex is recruited to chromatin as a single hexamer in the early G1 phase. Importantly, the cell cycles in these cells treated with Set8-siRNA for 26 h in total were not arrested completely, and some cells, especially S phase cells, slowly proceeded to the next cell cycle stage. Thus, we used a longer Set8-siRNA treatment ($> 40$ h) to determine the state of MCM at the late G1 phase.

### Supplementary Figures

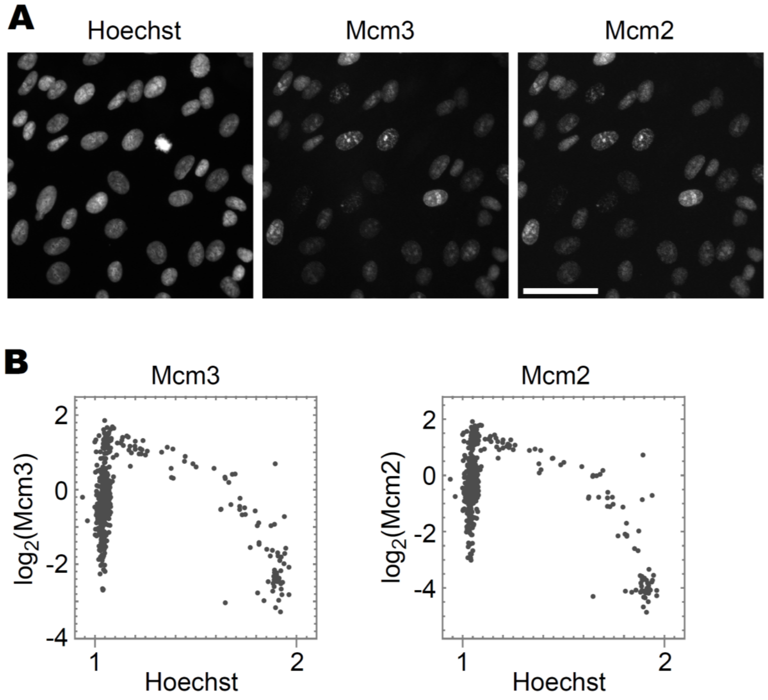


**Figure S1.** Single-cell plot analysis of chromatin-bound Mcm3 and Mcm2 proteins.

**A**. Representative fluorescence images of Mcm3 and Mcm2 in hTERT-RPE1 cells fixed by the pre-extraction method. Scale bar, 50 μm. **B**. Single-cell plot analysis based on the images in **A**. The number of cells examined in each panel is 400.

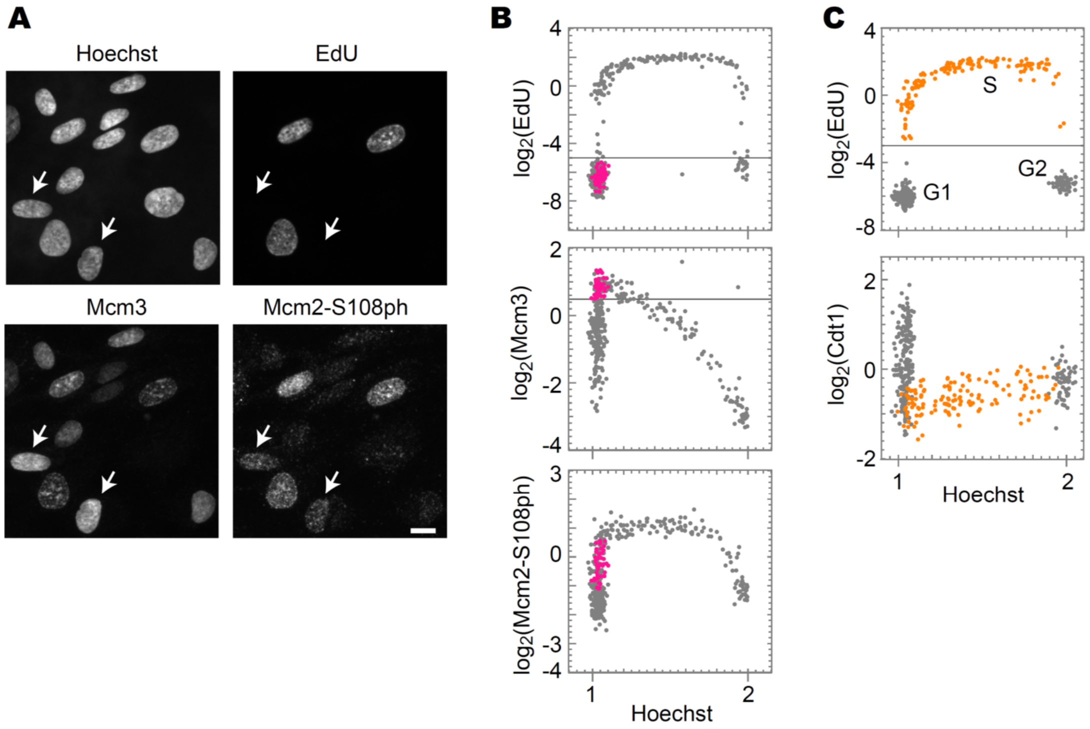


**Figure S2.** Single-cell plot analysis of phosphorylated Mcm2 and Cdt1.

**A**. Representative fluorescence images of Hoechst, EdU, Mcm3, and Mcm2-S108ph in hTERT-RPE1 cells prepared by the pre-extraction method. Arrows indicate representative cells with low EdU and high Mcm3 levels. Scale bar, 10 µm. **B.** Single-cell plot analysis of Mcm2-S108ph, Mcm3, and EdU based on the images in **A**. The G1 cells (Hoechst < 1.125 and $\log_2$(EdU) < –5) with high Mcm3 levels ($\log_2$(Mcm3) > 0.5) were selected and marked as the pink dots. The same cells were plotted in the Mcm2-S108ph panel. These cells marked with pink dots displayed relatively high levels of phosphorylation (–1 < $\log_2$(Mcm2-S108ph) < 0.5). The number of cells measured in each panel is 400. **C**. Single-cell plot analysis of EdU and Cdt1. The orange dots represent S phase cells based on EdU intensities ($\log_2$(EdU) > –3). The number of cells examined in each panel is 400.

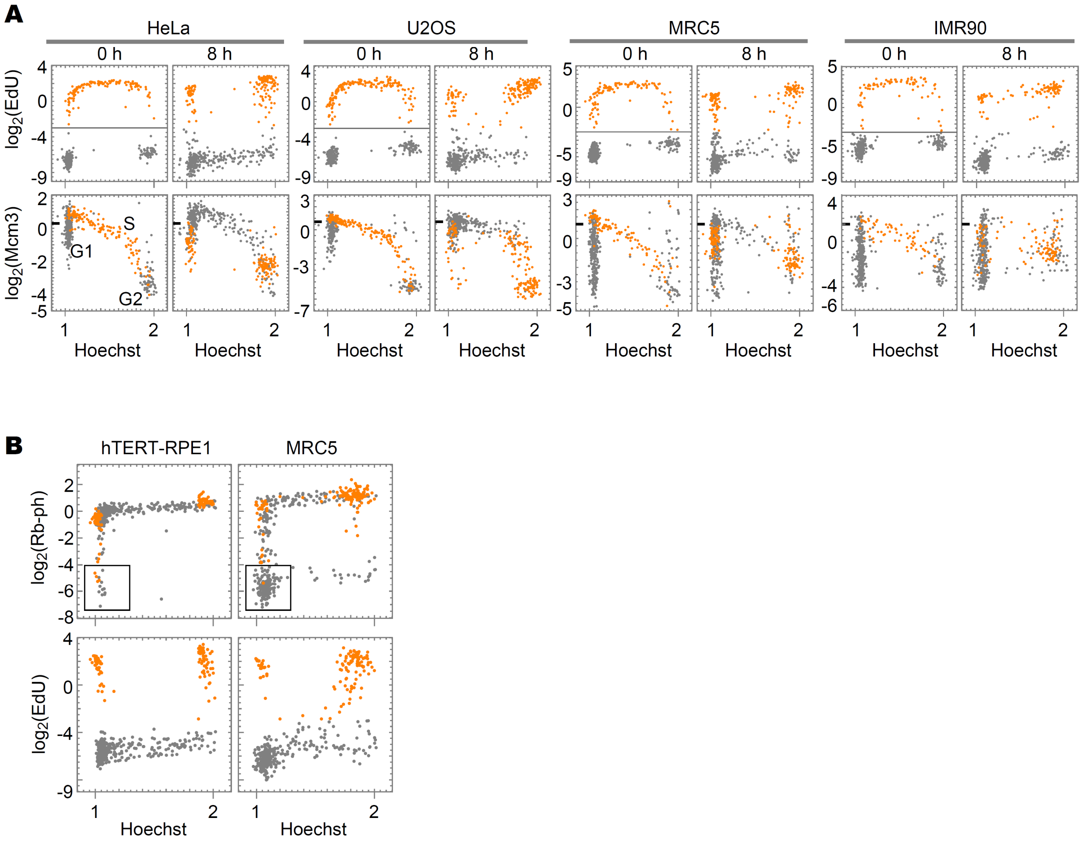


**Figure S3.** Single-cell plot analysis of chromatin-bound Mcm3 in various cell lines.

**A**. Single-cell plot analysis of Mcm3 and EdU in HeLa, U2OS, MRC5, and IMR90 cells in the growth phase. The cells were pulse-labeled with EdU for 30 min and chase-cultured without EdU. The cells were fixed at 0 or 8 h, as indicated, and applied to the single-cell plot analysis for EdU (upper panels) and Mcm3 (lower panels). The orange dots represent cells undergoing S phase at 0 h ($\log_2$(EdU) > –3). The number of cells examined in each panel is 450. The tick mark on the vertical axis of Mcm3 indicates a border between low and high levels. **B.** Single-cell plot analysis of phosphorylated Rb at Ser807/811 in hTERT-RPE1 and MRC5 cells. The cells were pulse-labeled with EdU for 30 min, and chase-cultured for 8 h. Then the cells were fixed by the direct fixation method. Orange dots represent cells undergoing S phase at 0 h ($\log_2$(EdU) > –3). The square regions indicate G1 cells with low levels of Rb phosphorylation. The number of cells examined in each panel is 450.

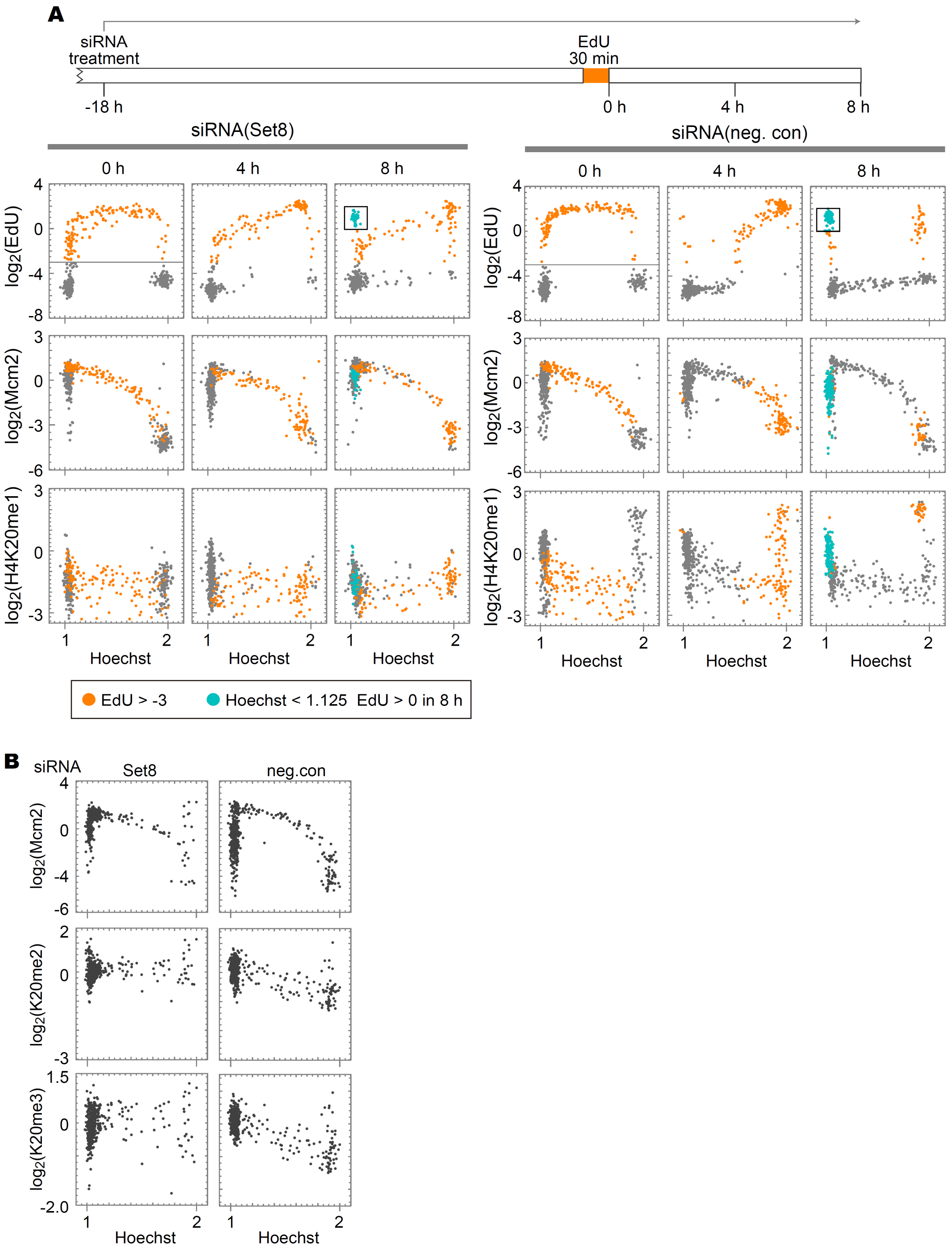


**Figure S4.** The effect of Set8-siRNA treatment on chromatin-bound Mcm2.

**A.** The cells were treated with Set8-siRNA or control-siRNA for 18 h and then pulse-labeled with EdU for 30 min, and chase-cultured in the presence of siRNAs until the indicated times (timescale on the top) and applied to the single-cell plot analysis (lower panels). The orange dots represent S phase cells at 0 h ($log_2$(EdU) > –3). The blue dots at 8 h represent the cells newly entering the G1 phase (Hoechst < 1.125, $log_2$(EdU) > 0, the squares in the EdU panels). The number of cells examined in each panel is 450. **B**. Single-cell plot analysis of Mcm2, H4K20me2, and H4K20me3 in cells treated with either Set8-siRNA or control-siRNA. The number of cells examined in each panel is 450.

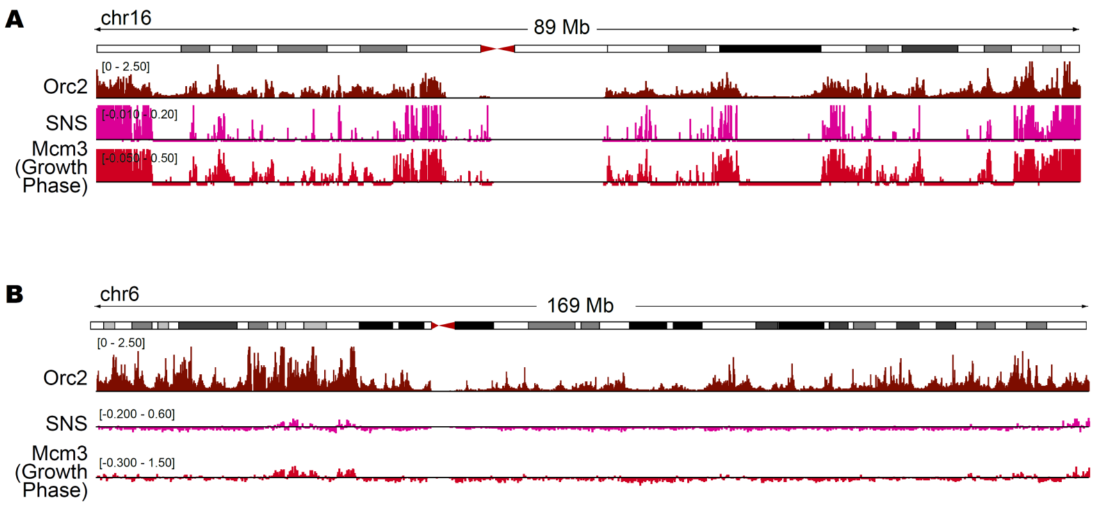


**Figure S5.** Genome-wide analysis of Mcm3 localization characterized using ChIP-seq analysis. Enrichment of Mcm3 (this study), Orc2 (K562 cells; GSM1717888), and SNS (IMR90 cells; GSM927235) are shown for chromosome 16 (**A**) and 6 (**B**). Vertical axes are shown on the linear scale for Orc2 (RPKM_ChIP), while the $\log_2$ scale was used for the others (RPKM_ChIP/RPKM_Input). The vertical axes for SNS-seq and Mcm3 in **A** are enlarged from those in Fig. 4C, whereas the vertical axes in **B** are shown in the same scale as in Fig. 4C.

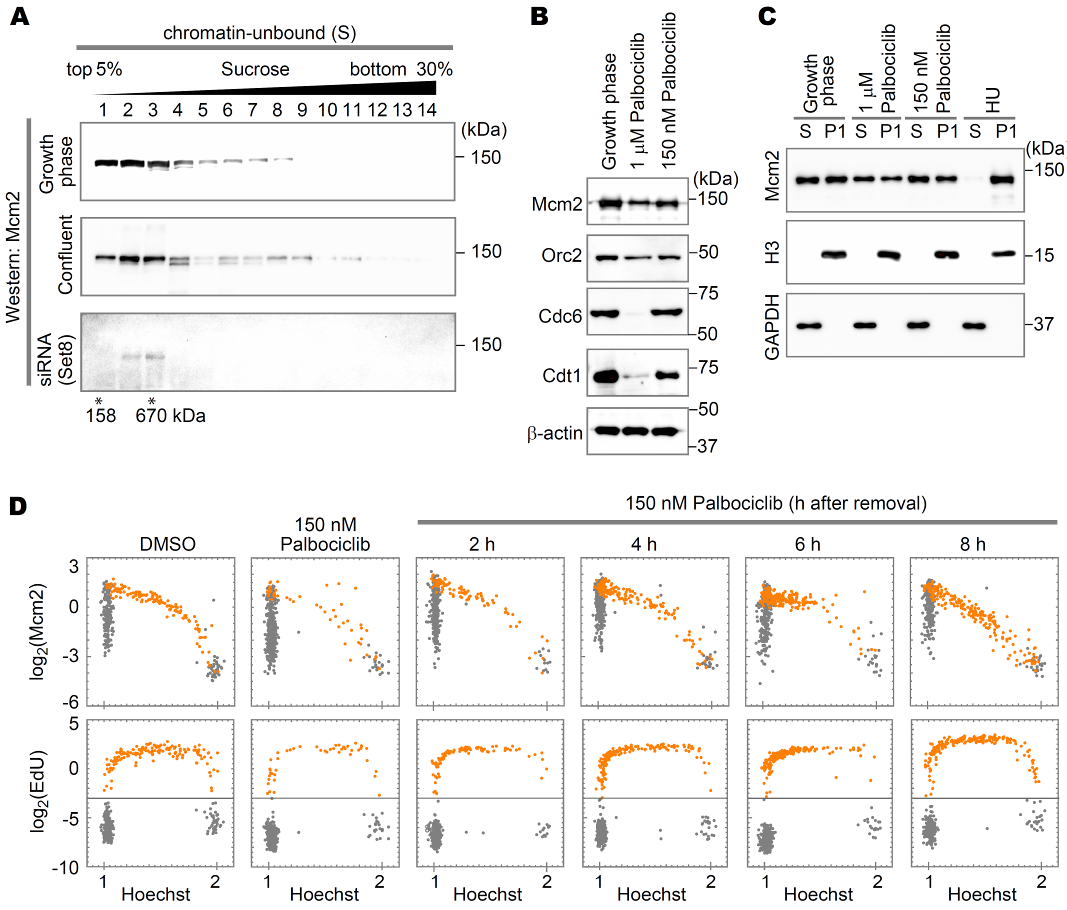


**Figure S6.** Characterization of MCM complexes in cell cycle-arrested cells

**A.** Western blot analysis of sucrose gradient fractions using anti-Mcm2 antibody. Chromatin-unbound (S) fraction was prepared from the growth phase cells, confluent cells, and Set8-siRNA-treated cells as described in Fig. 5A and fractionated through a linear 5–30% sucrose gradient. Molecular sizes derived from the sucrose gradient are indicated below the blots.

**B**. Western blot analysis using antibodies against Mcm2, Orc2, Cdc6, Cdt1, and β-actin. Whole cell lysate prepared from growth phase cells and palbociclib-treated cells (1 μM or 150 nM). The cells ($6 \times 10^5$ cells) were washed with PBS three times and then collected with 2× SDS loading buffer, followed by Western blot analysis. **C.** Western blot analysis of P1 and S fractions using antibodies against Mcm2, histone H3, and GAPDH. The fractions prepared from the growth phase cells, palbociclib-treated cells (1 μM or 150 nM), and 200 μM HU-treated cells.

**D**. Single-cell plot analysis of Mcm2 and EdU. Cells were arrested by 150 nM palbociclib for 18 h. The cells were released from the arrest by washing with pre-warmed culture medium. The cells were then cultured until collection at the indicated times and labeled with EdU for 30 min immediately before the collection. The orange dots represent S phase cells based on EdU intensities ($\log_2$(EdU) > –3). The number of cells examined in each panel is 450.

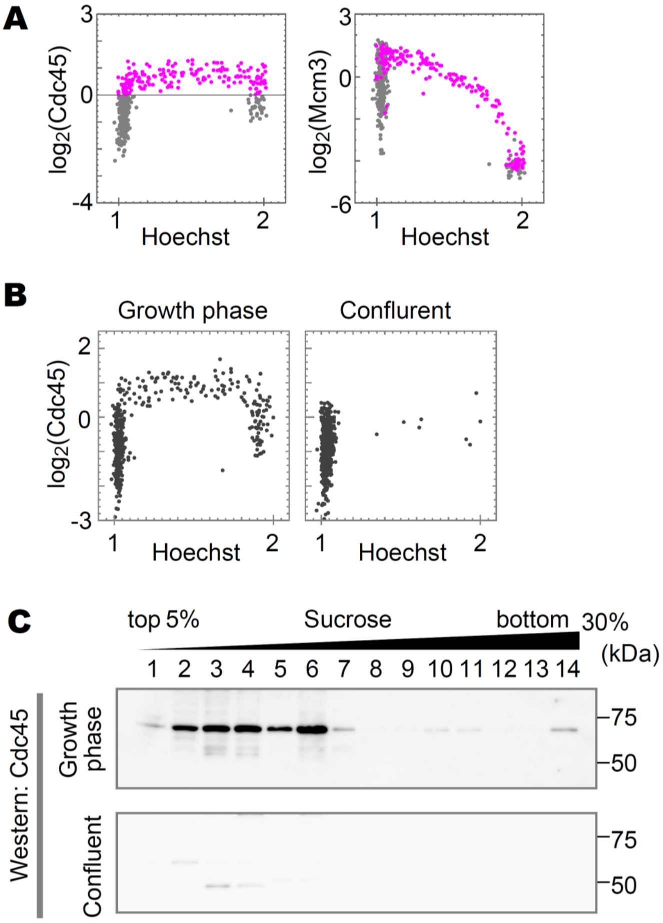


**Figure S7.** Cdc45 in single-cell plot analysis and sucrose gradient fractionation.

**A**. Single-cell plot analysis of Cdc45 and Mcm3 in hTERT-RPE1 cells in the growth phase. The pink dots represent the cells with high levels of Cdc45 ($\log_2$(Cdc45) > 0). The number of cells examined in each panel is 450. **B**. Single-cell plot analysis of Cdc45 in the growth phase, and confluent. The number of cells examined in each panel is 400. **C**. Western blot analysis of sucrose gradient fractions using anti-Cdc45 antibody. The P2+B fractions are the same preparations as for Fig. 5C.

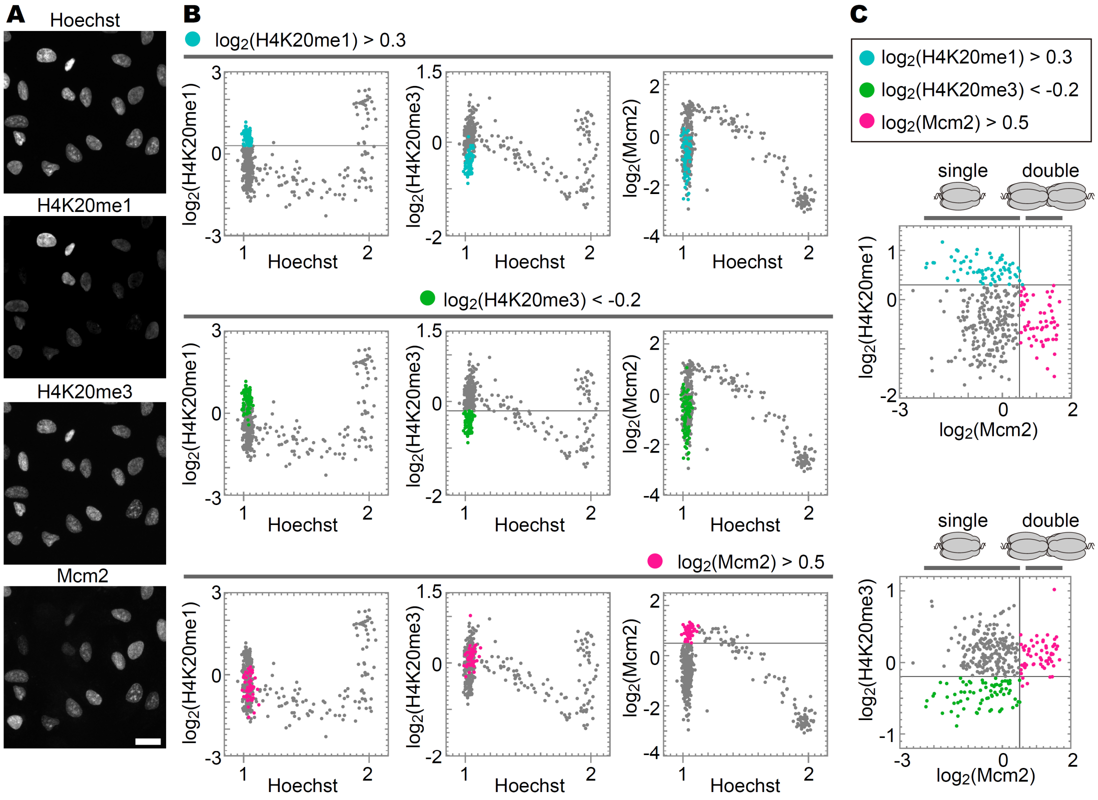


**Figure S8.** Relationship between histone H4K20 methylation levels and MCM hexamer states. **A.** Representative microscopic images of Hoechst, H4K20me1, H4K20me3, and Mcm2 in hTERT-RPE1 fixed by the pre-extraction method. Scale bar, 20 µm. **B.** Single-cell plot analysis of H4K20me1, H4K20me3, and Mcm2. In the top row, the G1 cells (Hoechst < 1.125) with high levels of H4K20me1 ($log_2$(H4K20me1) > 0.3) are shown in light blue; in the middle row, the G1 cells (Hoechst < 1.125) with low levels of H4K20me3 ($log_2$(H4K20me3) < –0.2) in green; in the bottom row, the G1 cells (Hoechst < 1.125) with high levels of Mcm2 ($log_2$(Mcm2) > 0.5) in pink. The number of cells examined in each panel is 450. **C**. Scatterplots between H4K20me1 and Mcm2 intensities, and between H4K20me3 and Mcm2 intensities based on **B**.

**Table S1**

| | Processed | Failed to align | | Suppressed due to –m | | Mapped | |
|---|---|---|---|---|---|---|---|
| | Reads | Reads | (%) | Reads | (%) | Reads | (%) |
| Growth Phase_MCM3IP | 22,386,500 | 2,902,003 | 12.96 | 6,174,348 | 27.58 | 13,310,149 | 59.46 |
| Growth Phase_Input | 20,199,698 | 426,817 | 2.11 | 4,395,194 | 21.76 | 15,377,687 | 76.13 |
| Confluent_MCM3IP | 24,168,260 | 3,428,799 | 14.19 | 6,773,629 | 28.03 | 13,965,832 | 57.79 |
| Confluent_Input | 20,602,678 | 424,843 | 2.06 | 4,671,977 | 22.68 | 15,505,858 | 75.26 |
| siRNA_KMT5A_MCM3IP | 29,458,846 | 3,144,117 | 10.67 | 8,326,223 | 28.26 | 17,988,506 | 61.06 |
| siRNA_KMT5A_Input | 26,531,902 | 543,461 | 2.05 | 5,500,653 | 20.73 | 20,487,788 | 77.22 |
